# Recipient-driven metabolic and epigenetic remodeling of donor hematopoietic cells after transplantation in Fanconi Anemia

**DOI:** 10.64898/2026.10.26.761890

**Authors:** Matilde Balbi, Stefano Regis, Giovanni Fiorito, Paolo Uva, Marta Maragliano, Fabio Corsolini, Federica Raggi, Luca Arcuri, Maria Carla Giarratana, Sara Pestarino, Ramona Tallone, Andrea Beccaria, Vanessa Cossu, Giorgia Damonte, Martina Serra, Grazia Maria Spaggiari, Carlo Dufour, Maura Faraci, Silvia Ravera, Filomena Pierri, Enrico Cappelli

## Abstract

**Background:** Haematopoietic cell transplantation (HCT) restores donor-derived haematopoiesis in Fanconi anaemia (FA) but leaves recipient stromal and other non-haematopoietic cells genetically uncorrected. We investigated to what extent the persistent FA recipient environment influences the metabolic, redox, inflammatory and epigenetic phenotype of donor-derived haematopoietic cells after HCT.

**Methods:** Peripheral blood and, when available, bone marrow samples were analysed from healthy donors, patients with FA before and after HCT, and severe aplastic anaemia (SAA) patients after HCT as transplant controls. Mitochondrial function, cellular energy status, oxidative damage, antioxidant responses and circulating cytokines were assessed. Healthy donor mononuclear cells were exposed to conditioned media from FANCA-deficient mesenchymal stromal cells or FA fibroblasts, with FANCA-corrected fibroblasts as controls. DNA methylation profiling, acute inflammatory stimulation and pharmacological targeting of inflammatory pathways or mitochondrial oxidative stress were also performed.

**Results:** Despite stable full donor chimerism, FA recipients retained a pro-inflammatory and pro-oxidative circulating environment after HCT that was not observed in SAA transplant controls. Donor-derived mononuclear cells exposed to this environment acquired a distinct dysfunctional phenotype characterized by impaired mitochondrial oxidative phosphorylation and energy homeostasis, compensatory antioxidant and stress-response activation, and persistent lipid and protein oxidation. These alterations remained detectable after 96 h of ex vivo culture, suggesting relatively stable cellular remodelling, and were accompanied by a DNA methylation profile that, although globally shifted toward the donor state, retained an intermediate FA-associated episignature. Mechanistically, conditioned media from FANCA-deficient stromal cells and FA fibroblasts reproduced mitochondrial and oxidative abnormalities in healthy donor cells, whereas FANCA-corrected fibroblasts did not. Pharmacological targeting of inflammatory pathways or mitochondrial oxidative stress improved energy homeostasis and reduced oxidative damage, supporting a contributory role for chronic inflammation and oxidative stress in the microenvironment-driven phenotype acquired by donor-derived cells.

**Conclusions:** HCT corrects FA haematopoiesis without fully normalising the recipient environment. Persistent exposure to the FA milieu promotes metabolic, redox, signalling and epigenetic remodelling of donor-derived haematopoietic cells after transplantation. These findings identify the recipient environment as a determinant of post-HCT cell biology and highlight chronic inflammation and oxidative stress as mechanisms contributing to donor-cell remodelling in FA.

## 1. BACKGROUND

Fanconi Anemia (FA) is an inherited BM failure syndrome caused by defects in the FA/BRCA DNA repair pathway [1–3]. The hallmark FA clinical manifestation is progressive BM failure[1,4]. In addition to defective DNA repair, FA cells exhibit mitochondrial dysfunction, increased oxidative stress, and impaired antioxidant defences [5–7].

Direct experimental evidence further suggests that the FA metabolic and redox phenotype is largely unmasked when cells leave the hypoxic BM niche for more oxygenated conditions. Specifically, FA BM mononuclear cells (MNCs) display minimal metabolic abnormalities under hypoxia but, under normoxia, develop altered oxidative phosphorylation, increased oxidative stress, impaired activation of endogenous antioxidant defences, and G2/M cell-cycle arrest [6–8]. In addition, Inflammatory and stress-signalling mechanisms play a central role, as FA-HPCs are hypersensitive to oxidative stress and apoptotic cytokines such as TNF-α and IFN-γ [1,9]. Chronic type-I interferon signalling has been proposed as one inflammatory axis linking genome instability to premature aging and marrow pathology in FA, and recent studies support elevated cGAS-STING activity in FANCA- and FANCD2-deficient cells [10,11]. In parallel, hyperactive TGF-β signalling directly contributes to marrow failure, as TGF-β inhibition improves FA cell survival and rescues HSPC proliferative and functional defects in mouse and patient-derived models [4]. Collectively, these findings support the multifactorial view of BM failure in FA, arising from the convergence of genomic instability, mitochondrial dysfunction, oxidative stress, and chronic inflammatory signalling [8,12,13]. HSC transplantation (HCT) remains the only curative treatment for FA haematological manifestations [14–16] with remarkable improvements in outcomes over recent decades [14,16–19]. However, transplantation does not correct the genetic defect in non-hematopoietic recipient tissues, including stromal and niche-associated cells that are involved in HSC quiescence, self-renewal, differentiation and survival [20–27]

Chimerism analyses in transplanted FA patients demonstrated that BM MSCs remain entirely of recipient origin following conditioning, indicating that this compartment is not replaced by transplantation[3]and that the FA microenvironment retains FA-related biological features, thus raising the important, so far unsolved question of whether the FA microenvironment continues to impose inflammatory, oxidative, metabolic, and epigenetic stress on donor-derived hematopoietic cells after HCT. Evidence from the literature on FA and inherited marrow-failure supports the biological plausibility of this hypothesis, showing a chronic pro-inflammatory cytokine milieu involving TNF-α, IL-6, TGF-β, IFN signalling, and related microenvironment effects [28,29]. This question has important clinical implications because, despite successful HCT, patients with FA remain at substantial risk of long-term complications, including the increased risk of malignancies as compared with non-transplanted FA patients [12,16,17].

In this study, we investigated whether the FA hematopoietic microenvironment remains pro-oxidative and pro-inflammatory after HCT and whether this persistent exposure of donor-derived hematopoietic cells may drive their long-term metabolic and epigenetic remodelling.

## 2. METHODS

### 2.1 Reagents

Unless otherwise specified, analytical-grade reagents were obtained from Sigma-Aldrich (St. Louis, MO, USA). Ultrapure water was used for all biochemical assays.

### 2.2 Human samples and ethical approval

Peripheral blood (PB) and, when available, bone marrow (BM) mononuclear cells (MNCs) were obtained from healthy donors (HDs), patients with Fanconi anaemia (FA), FA patients who had undergone haematopoietic cell transplantation (FA post-HCT), and patients with severe aplastic anaemia who had undergone HCT (SAA post-HCT),all recruited at the Hematology Unit of the IRCCS Istituto Giannina Gaslini. The SAA post-HCT group served as a disease-control population to assess whether the alterations observed in FA post-HCT patients were specifically associated with the underlying FA condition rather than representing a general consequence of HCT.

Samples were collected after written informed consent from all participants or legal guardians, in accordance with the Declaration of Helsinki and approved by the local Ethics Committee under protocol JS 002, initially approved on 24 September 2010 and most recently re-approved after amendments on 21 April 2026.

HD plasma was separated by centrifugation at 3,000 g for 10 minutes, at 25°C, aliquoted, and stored at-80 °C until analysis.

For a subset of transplanted patients, the corresponding healthy stem cell donors were also studied. For all transplanted patients, demographic and transplant-related characteristics were collected, including age, sex, donor type, hematopoietic stem cell transplantation (HCT) platform, and time from HCT to sample collection. For patients with FA, the indication for HCT and the FA complementation group were also recorded.

### 2.3 Isolation and culture of BM and PB-MNCs

PB-and BM-MNCs were isolated by Ficoll-Histopaque density-gradient centrifugation, as described in [6]. MNCs were cultured in RPMI-1640 medium (GIBCO, Billing, MT, USA) supplemented with 10% fetal bovine serum (FBS) (EuroClone, Milan, Italy) and penicillin/streptomycin (EuroClone, Milan, Italy) at 37 °C in a humidified atmosphere containing 5% CO₂. For persistence experiments, freshly isolated PB-MNCs were maintained ex vivo for 96 h under identical culture conditions before biochemical, oxidative, and metabolic analyses.

### 2.4 Culture of healthy, FANCA-silenced MSCs and FA MSCs

Healthy donor mesenchymal stromal cells (HDMSCs), scramble MSCs (scrMSCs), FANCA gene-silenced MSCs (shFAMSCs), and MSCs derived from FA patients (FA MSCs), all derived from BM, were cultured in MesenPRO RS™ Medium (#12746-012, Thermo Fisher Scientific, Waltham, MA, USA) with 2 mM glutamine and penicillin/streptomycin (EuroClone, Milan, Italy) at 37°C and 5% CO₂. When necessary, MSCs were cultured with plasma derived from HD for 72 h.

FANCA gene silencing was performed using specific siRNAs (#4392420, Assay IDs s162, s163, s164, Thermo Fisher Scientific, Waltham, MA, USA). Scramble-transfected MSCs (#4390843, Thermo Fisher Scientific, Waltham, MA, USA) were used as negative controls. FANCA knockdown efficiency was verified by real-time PCR using FANCA-and GAPDH-specific primers and probes (4448892, Assay ID Hs01116668_m1, and #4351370, Assay ID Hs99999905_m1, Thermo Fisher Scientific, Waltham, MA, USA).

### 2.5 Culture of FA and FA-corrected fibroblasts

Primary FANCA-mutated fibroblasts (FA fibroblasts) and isogenic FANCA-corrected fibroblasts (FA-corr fibroblasts) derived from six patients were obtained from the “Cell Line and DNA Biobank from patients affected by Genetic Diseases” (G. Gaslini Institute)—Telethon Genetic Biobank Network (Project No. GTB07001). FA-corr controls were generated by correction with the S11FAIN retroviral vector. Primary fibroblasts were cultured as a monolayer in DMEM high glucose with glutamax ® (GIBCO, Billing, MT, USA), containing 10% FBS (EuroClone, Milano, Italy), and penicillin/streptomycin (EuroClone, Milan, Italy) at 37°C with5% CO_2_. When necessary, fibroblasts were cultured with plasma derived from HD for 72 h.

### 2.6 Preparation of conditioned media and exposure of HD PB-MNCs

Conditioned media were generated from HDMSCs, scrMSCs, shFAMSCs, FA MSCs, FA fibroblasts, and FA-corr fibroblasts. Briefly, cells were washed with PBS and incubated in fresh medium, as specified above, for 24 h. Supernatants were collected, centrifuged at 3,000 g for 10 minutes to remove cells and debris, and used immediately.

HD PB-MNCs were cultured either in standard medium (as reported in section 2.3) or in conditioned media derived from the indicated MSCs or fibroblast cultures for 24 h.

### 2.7 Pharmacological stimulation and pathway inhibition

To model acute inflammatory or infection-like stimulation, PB-MNCs were treated with R848/resiquimod, a Toll-like receptor 7/8 agonist, at 1 µM for 24 h, according to previously described stimulation conditions in FA cellular models [30].

For rescue experiments, HD PB-MNCs and FA post-HCT PB-MNCs were treated with inhibitors targeting TNF-α, TGF-β, and NLRP3-related inflammatory pathways, or with the mitochondria-targeted antioxidant MitoQ. Cells were treated for 24 h with the following compounds: 10 μg/mL etanercept (TNF-α inhibition), 10 μg/mL luspatercept (modulation of TGF-β superfamily signalling), 10 μM MCC950 (NLRP3 pathway inhibition), or 100 μM MitoQ (mitochondria-targeted antioxidant activity). All compounds were obtained from Sigma-Aldrich (St. Louis, MO, USA). For combined stimulation experiments, cells were exposed to R848 alone or together with the indicated inhibitor/antioxidant. Vehicle-treated cells were included as controls.

### 2.8 Biochemical, molecular and epigenetic analyses

Full experimental procedures, reagent details and analytical parameters are provided in the Supplementary Methods.

Extracellular vesicles (EVs) were isolated from 500 μl of HD and FA plasma using the exoRNeasy Serum/Plasma Midi Kit (Qiagen), following pre-clearing centrifugation and the manufacturer’s protocol [31].

Cell homogenates were prepared by sonication in PBS containing protease and phosphatase inhibitors, and protein concentrations were determined by Bradford assay.

Oxidative phosphorylation was assessed in digitonin-permeabilized PB-MNCs or isolated EVs by measuring ATP synthesis through a luciferin–luciferase assay and oxygen consumption using an amperometric electrode, with pyruvate/malate or succinate as respiratory substrates; coupling efficiency was expressed as the P/O ratio [6,32].

Cellular ATP and AMP were quantified by enzyme-coupled spectrophotometric assays and expressed as the ATP/AMP ratio [6].

Oxidative damage was evaluated by measuring MDA through the TBARS assay and 4-HNE, nitrotyrosine and 8-OHdG using commercial ELISA kits, whereas glutathione reductase and catalase activities were determined spectrophotometrically [33].

Plasma TGF-β1, TNF-α, IL-6 and IL-8 were quantified using Simple Plex assays on the Ella platform. NRF2, phosphorylated and total STAT2, and phosphorylated and total SMAD3 were analysed by Western blot, with densitometric values normalized to actin.

Genome-wide DNA methylation profiling of PBMC DNA was performed using the Illumina Methylation Screening Array. Data were pre-processed with the sesame R package, corrected for batch effects using ComBat and analysed by principal component analysis; methylation-derived scores for 109 circulating proteins were calculated using MethylDetectR [34–37].

### 2.9 Statistical analysis

Data were analysed using Prism 10 Software. Results are reported as mean ± standard deviation (SD). Normality was assessed using the Shapiro–Wilk test when appropriate. Comparisons between two groups were performed using unpaired or paired Student’s t-test according to the experimental design. Multiple-group comparisons were performed using one-way analysis of variance (ANOVA), followed by Tukey’s post hoc test. Differences were considered statistically significant at p < 0.05. The number of biological replicates and independent experiments is reported in each figure legend.

## 3. RESULTS

### 3.1 Patient population

The study included 15 FANCA patients and 15 patients with SAA. The characteristics of the study population are summarized in Table 1. In 14/15 FA patients, the indication for HCT was bone marrow failure, and it was a poor-risk SF3B1 mutation. At the time of analysis, 13 of 15 patients with FA (86.7%) and all 15 patients with SAA (100%) were alive. Stable full donor chimerism was observed in all patients included in the study. No significant differences were found between the two groups apart from time from HCT to sample collection, which was significantly longer in patients with FA than in those with SAA (3.78 ± 3.33 vs 1.78 ± 1.82 years, respectively; p = 0.037). There was a statistically significant difference in donor age between the FA and SAA groups (34.38 ± 16.15 vs. 38.71 ± 12.72 years, *p*< 0.001). The haploidentical donor for each patient with FA was one of the parents and, in every case, was a carrier of a FANCA-associated pathogenic variant.

**Table 1.** FA and SAA patients and transplant characteristics: MSD (Matched Sibling donor), MUD (Matched Unrelated Donor), Haplo (Haploidentical donor), HCT (Hematopoietic Stem Cell Transplantation), TCD (T cell depletion), PTCy (post-transplant cyclophosphamide)

|  | FA patients | SAA patients | p |
| --- | --- | --- | --- |
| **N** | 15 | 15 |  |
| **M/F** | 11/4 | 10/5 | 0,093 |
| **Age (y) at pre HCT sampling (Mean (sd))** | 11,39 (5,85) | 11,20 (7,47) | ns |
| **Donor Type** | MUD 6/15 (40%) MSD 3/15 (20%) Haplo 6/15 (40%) all ex vivo TCD | MUD 7/15 (46,7%) MSD 2/15 (13,3%) Haplo 6/15 (40%), all ex vivo TCD 5PTCy | ns |
| **Age (y) at post HCT sampling (Mean ± SD)** | 15,17 ± 5,98 | 12,97 ± 7,61 | ns |
| **Time from HCT at sampling** | 3,78 ± 3,33 | 1,78 ± 1,82 | 0,037 |
| **Alive n° (%)** | 13/15 (86,6%) | 15/15 (100%) | ns |

### 3.2 FA plasma is characterized by increased lipid and protein oxidation

To determine whether the hematopoietic environment of FA patients is characterized by oxidative stress, markers of lipid and protein oxidation in BM and PB plasma have been quantified. No significant difference in MDA content was observed between BM and PB plasma from HD. Conversely, a significant increase in MDA was evident in both BM and PB plasma from non-transplanted FA patients compared with HD. Interestingly, FA PB plasma showed higher MDA levels than FA BM plasma, indicating that oxidative stress is maintained, and possibly amplified, in the peripheral compartment (Fig. 1A).

**Figure 1.**
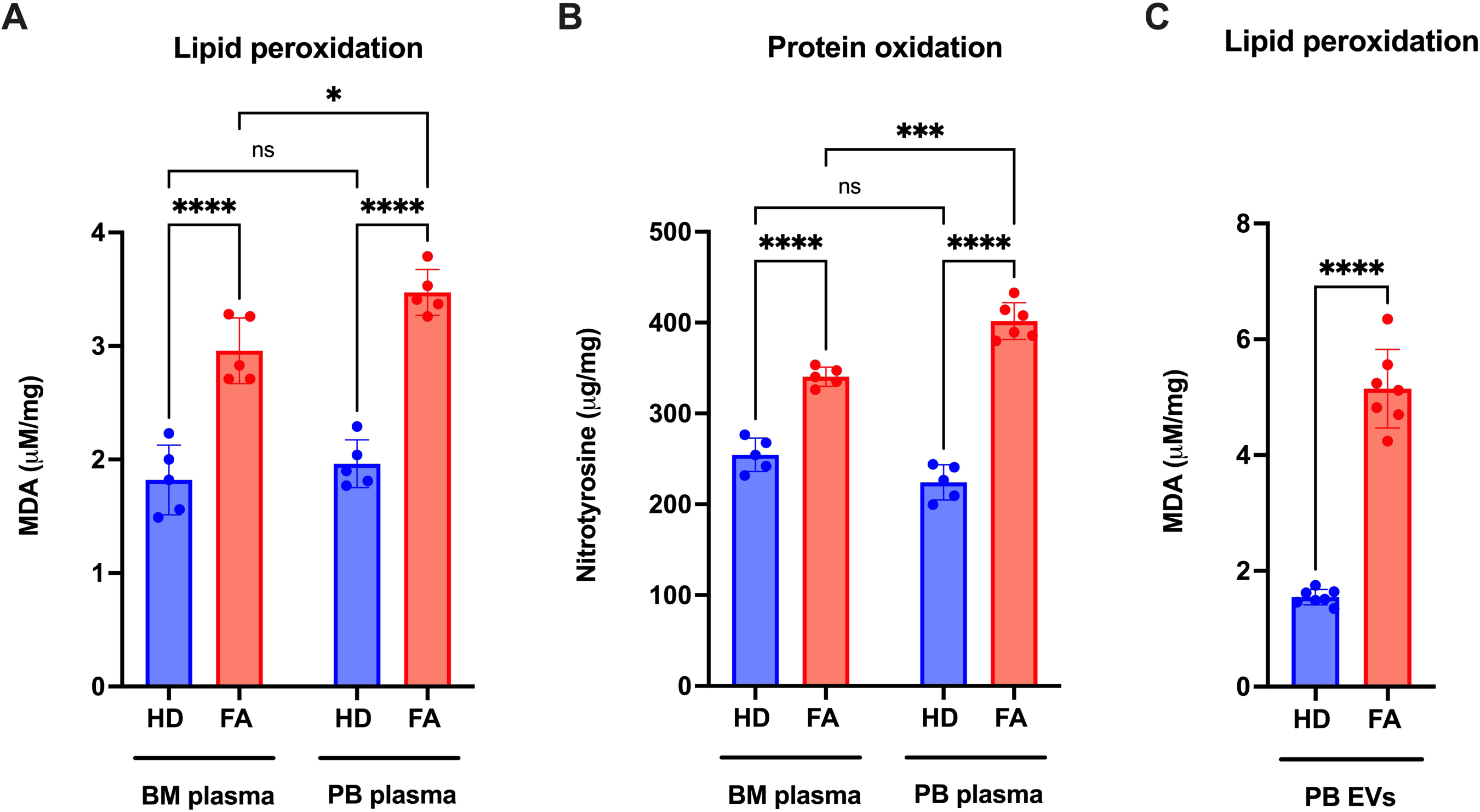
Assessment of oxidative damage markers in plasma and extracellular vesicles. (A) Lipid peroxidation measured as malondialdehyde (MDA) levels in bone marrow (BM) and peripheral blood (PB) plasma from healthy donors (HD) and non-transplanted patients with Fanconi Anemia (FA). (B) Protein oxidation assessed by nitrotyrosine levels in BM and PB plasma from HD and FA subjects. (C) MDA levels in extracellular vesicles isolated from peripheral blood (PBEVs) of HD and FA subjects. Data are presented as mean ± SD (n= 5 for (A) and (B); n= 7 for (C)). Statistical analysis was performed using one-way ANOVA, followed by Tukey’s post hoc multiple comparison test for (A) and (B), and paired Student’s t-test for (C). ns, not significant; *p < 0.05; ***p < 0.001; ****p < 0.0001.

A similar pattern was observed for protein oxidation. Nitrotyrosine levels were significantly higher in FA BM and PB plasma than in the corresponding HD. As observed for MDA, nitrotyrosine levels were comparable between HD BM and PB plasma, whereas FA PB plasma displayed higher levels than FA BM plasma (Fig. 1B). In addition, extracellular vesicles isolated from FA PB plasma showed markedly increased MDA levels compared with HD PB-EVs, indicating that circulating EVs also carry a pro-oxidative signature in FA patients (Fig. 1C).

Together, these data indicate that the FA hematopoietic environment is characterized by increased lipid and protein oxidation both in the BM and PB compartments, with oxidative damage also detectable in circulating extracellular vesicles.

### 3.3 FA mesenchymal stromal cells display mitochondrial dysfunction, impaired OxPhos efficiency, and increased lipid peroxidation

To investigate whether FA-BM niche cells may participate in the generation of a pro-oxidative microenvironment, the bioenergetic and redox profile of MSCs silenced for the FANCA gene has been analysed. Compared with HD and scr MSCs, shFA MSCs and non-transplanted FA MSCs showed a profound alteration of mitochondrial OxPhos. In the presence of P/M, shFA MSCs and FA MSCs displayed a significant reduction in ATP synthesis and OCR, together with reduced OxPhos efficiency, expressed as P/O ratio (Fig. 2A–C). With Succ as substrate, shFA MSCs and FA MSCs showed increased ATP synthesis and respiratory activity compared with control cells, but this was accompanied by a reduced P/O ratio, indicating an inefficient and partially uncoupled mitochondrial phenotype (Fig. 2A–C).

**Figure 2.**
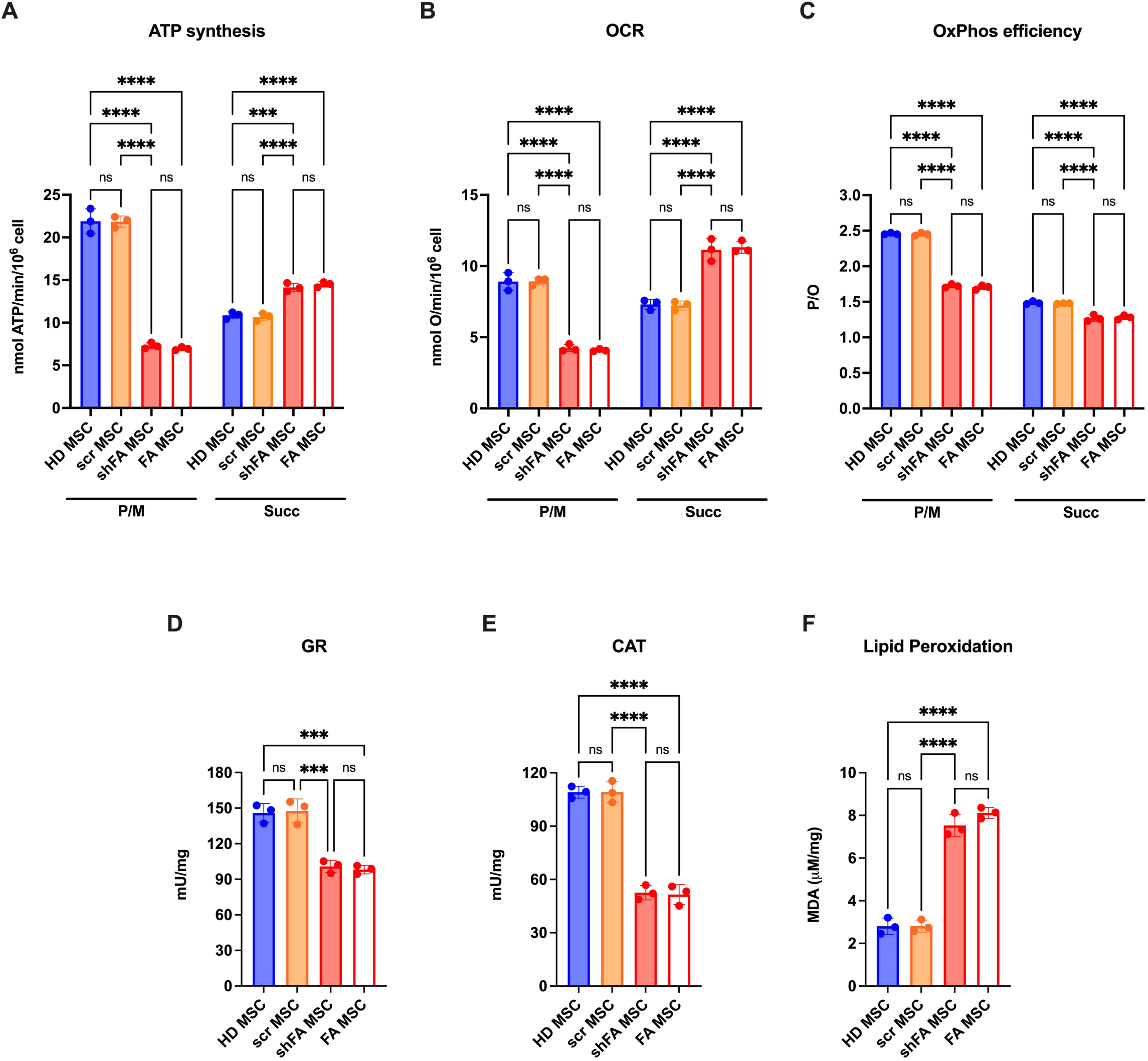
Bioenergetic, antioxidant defences, and oxidative damage parameters in MSCs. (A) ATP synthesis; (B) Oxygen consumption rate (OCR); (C) P/O ratio, calculated as the amount of ATP synthesized per oxygen atom consumed;(D) Glutathione reductase (GR) activity. (E) Catalase (CAT) activity; (F) Lipid peroxidation assessed by malondialdehyde (MDA) levels in healthy donor mesenchymal stromal cells (HD MSC), scrambled control mesenchymal stromal cells (scr MSC), Fanconi Anemia-silenced mesenchymal stromal cells (shFA MSC), and Fanconi Anemia mesenchymal stromal cells (FA MSC). For panels A-C, pyruvate plus malate (P/M) or succinate (Succ) were used as respiratory substrates. Data are presented as mean ± SD (n= 3 for each panel). Statistical analysis was performed using one-way ANOVA, followed by Tukey’s post hoc multiple comparison test. ns, not significant; ***p < 0.001; ****p < 0.0001.

Consistent with this mitochondrial defect, shFA MSCs and FA MSCs showed a reduced antioxidant enzymatic response, with significantly lower glutathione reductase and catalase activities compared with Wt MSCs (Fig. 2D and 2E). This impaired antioxidant profile was associated with a marked increase in MDA levels, indicating enhanced lipid peroxidation (Fig. 2F).

These results show that FA-gene mutated MSCs have a substrate-dependent mitochondrial dysfunction associated with reduced OxPhos efficiency, defective antioxidant defenses, and accumulation of lipid oxidative damage, confirming the altered mitochondrial profile previously observed in lymphocytes, lymphoblasts, and fibroblasts carrying the FANCA gene mutation [38–40].

### 3.4 FA MSCs and FA fibroblasts induce oxidative modification of HD PB plasma lipids and proteins

To evaluate whether FA MSCs can transfer oxidative stress to the extracellular environment, HD PB plasma was incubated on HD MSCs, scr MSCs, shFA MSCs, or FA MSCs, and oxidative damage markers were then measured in the plasma after culture.

HD PB plasma incubated with HD MSCs or scr MSCs did not show significant increases in MDA or nitrotyrosine levels. In contrast, HD PB plasma exposed to shFA MSCs or FA MSCs showed a marked increase in both lipid peroxidation and protein oxidation (Fig. 3A and 3B).

**Figure 3.**
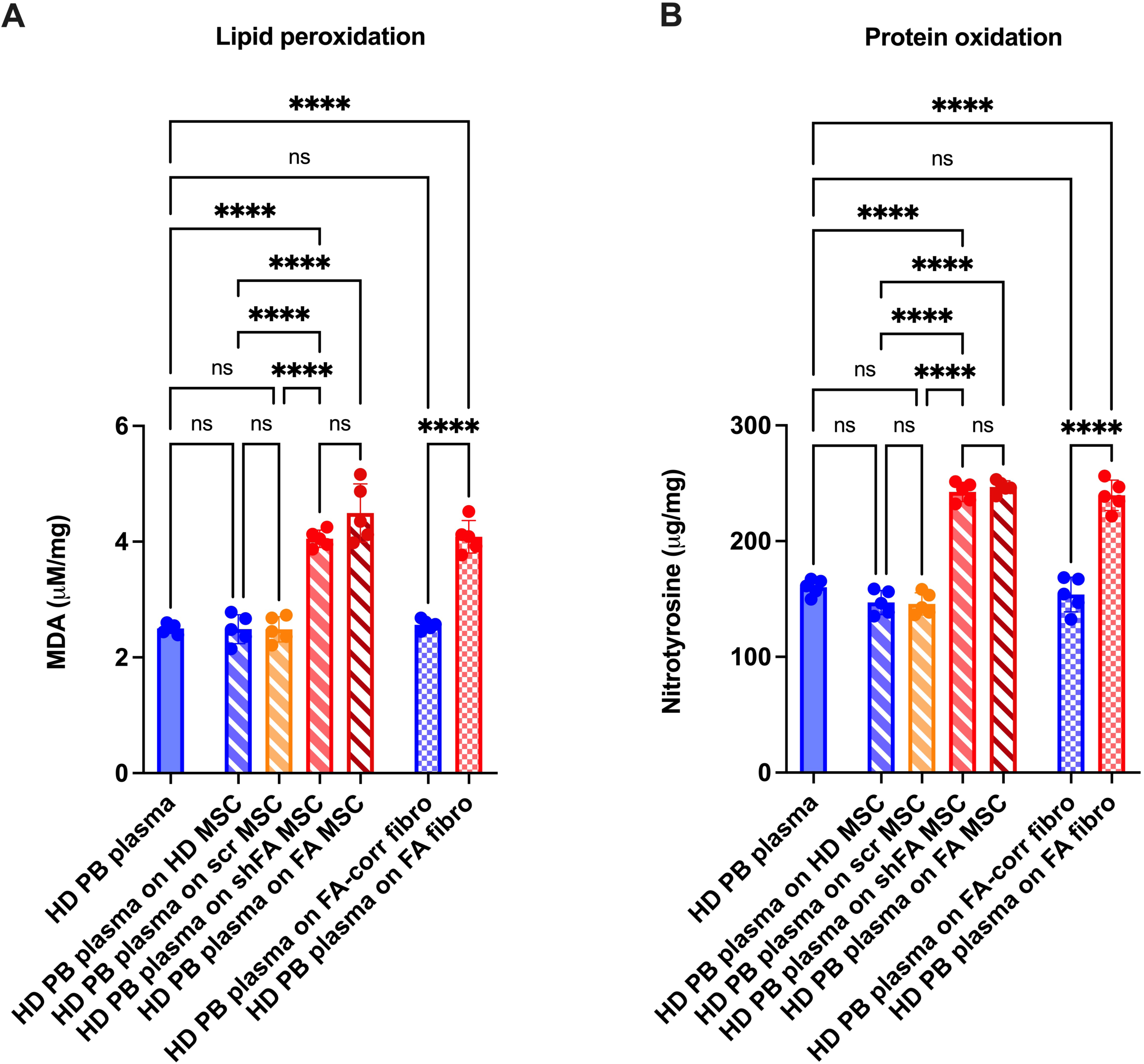
Oxidative damage in HD PB plasma added on HD, shFA, or FA MSCs. (A) MDA levels in healthy donor (HD) peripheral blood (PB) plasma, HD PB plasma incubated with healthy donor mesenchymal stromal cells (HD MSC), scrambled mesenchymal stromal cells (scr MSC), FANCA-silenced mesenchymal stromal cells (shFA MSC), Fanconi Anemia mesenchymal stromal cells (FA MSC), FANCA-corrected fibroblasts (FA-corr fibro), or FANCA-mutated fibroblasts (FA fibro). (B) Protein oxidation, assessed by nitrotyrosine levels, in HD plasma, HD plasma incubated with healthy MSC, scr MSC, shFA MSC, FA-corr fibro, or FA fibro. Data are presented as mean ± SD (n= 5 for each panel). Statistical analysis was performed using one-way ANOVA, followed by Tukey’s post hoc multiple comparison test. ns, not significant; ****p < 0.0001.

To determine whether this effect was not restricted to MSCs, HD PB plasma was also incubated with FA fibroblasts or FA-corr fibroblasts. Similarly, PB plasma exposed to FA fibroblasts, but not to FA-corr fibroblasts, showed significantly increased MDA and nitrotyrosine levels (Fig. 3A and 3B).

These findings indicate that FA MSCs and FA fibroblasts can promote oxidative modifications in the extracellular compartment, leading to oxidation of plasma components, including lipids and proteins.

### 3.5 FA MSCs-and FA fibroblast-conditioned environments impair mitochondrial metabolism and cellular energy status in HD MNCs

To determine whether the oxidative microenvironment generated by FA stromal cells has functional consequences on hematopoietic cells, HD PB-MNCs were cultured in media conditioned by controls (i.e., HD MSCs and scr MSCs) or shFA MSCs. HD PB-MNCs cultured in medium conditioned by both controls-maintained ATP synthesis, OCR, and OxPhos efficiency comparable to the untreated sample, both in the presence of P/M and Succ as respiratory substrates (Fig. 4A–F). In contrast, medium conditioned by shFA MSCs significantly impaired mitochondrial metabolism, reducing ATP synthesis, OCR, and P/O ratio under both complex I-linked and complex II-linked substrate conditions in HD PB-MNCs (Fig. 4A–F).

**Figure 4.**
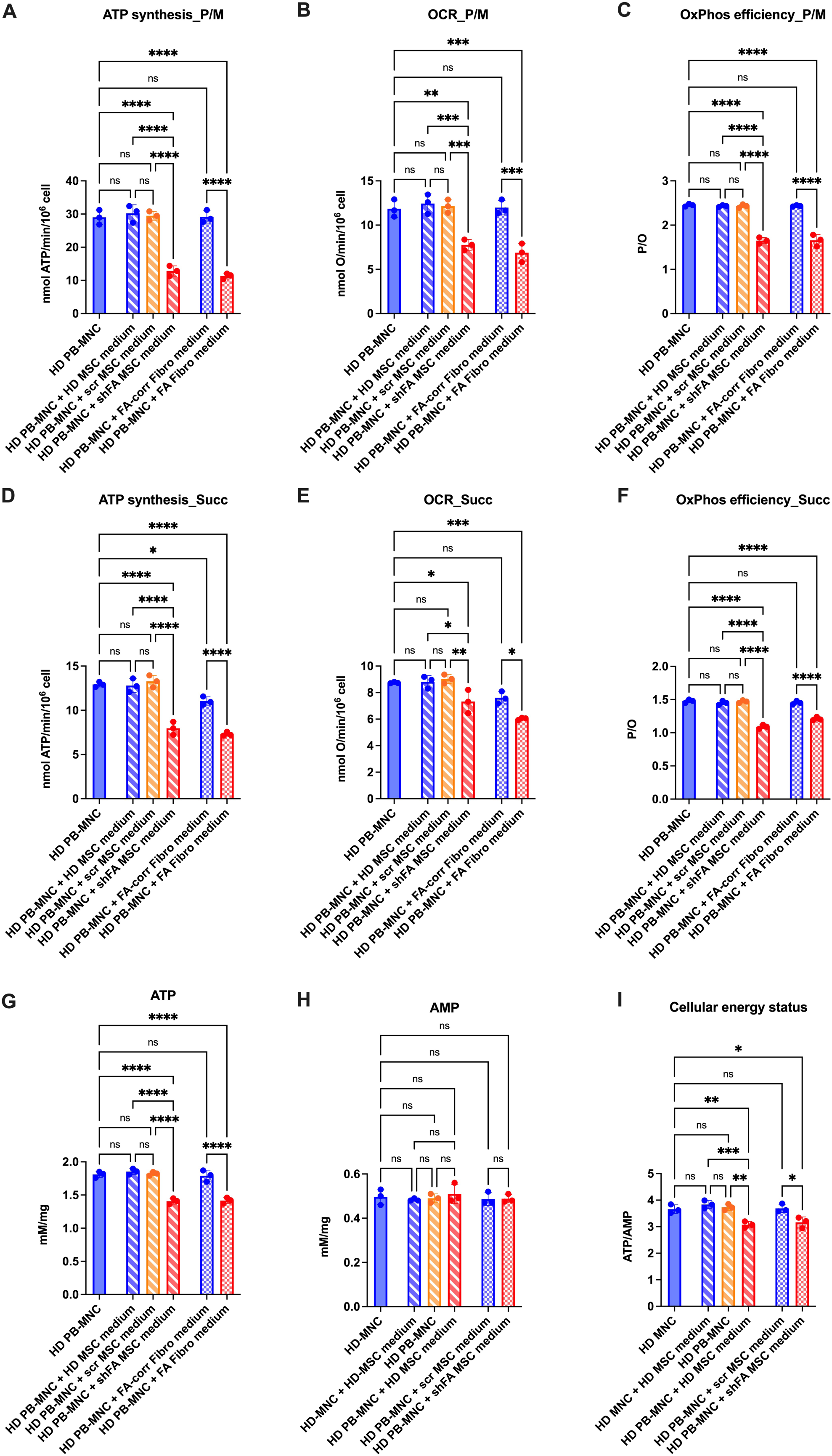
Mitochondrial metabolism and cellular energy status in HD-MNCs cultured with stromal-cell- and fibroblast-conditioned media. Healthy donor peripheral blood mononuclear cells (HD PB-MNCs) were cultured alone or in conditioned media derived from HD mesenchymal stromal cells (HD MSC medium), scramble-control MSCs (scr MSC medium), FANCA-silenced MSCs (shFA MSC medium), FANCA-corrected fibroblasts (FA-corr fibro medium) or FANCA-mutated fibroblasts (FA fibro medium). Mitochondrial ATP synthesis (A), oxygen consumption rate (OCR) (B) and OxPhos efficiency, expressed as P/O ratio (C), were measured using pyruvate/malate (P/M) as complex I-linked substrates. ATP synthesis (D), OCR (E) and P/O ratio (F) were also measured using succinate (Succ) as a complex II-linked substrate. Intracellular ATP (G), AMP (H) and cellular energy status, expressed as ATP/AMP ratio (I), were assessed in the same experimental conditions. Data are presented as mean ± SD (n= 3 for each panel). Statistical analysis was performed using one-way ANOVA, followed by Tukey’s post hoc multiple comparison test. ns, not significant; *p < 0.05; **p < 0.01; ***p < 0.001; ****p < 0.0001.

A similar mitochondrial defect was observed when HD PB-MNCs were exposed to medium conditioned by FA fibroblasts. FA fibroblast-conditioned medium significantly reduced ATP synthesis, OCR, and OxPhos efficiency compared with untreated HD PB-MNCs (Fig. 4A–F). Conversely, medium from FA-corr fibroblasts did not induce a comparable impairment, indicating that the metabolic defect was specifically associated with the FA cellular background.

Consistent with the reduced mitochondrial ATP-generating capacity, HD PB-MNCs exposed to shFA MSC- or FA fibroblast-conditioned media showed decreased intracellular ATP levels and stable AMP levels (Fig. 4G and 4H). This resulted in a reduction of the ATP/AMP ratio, indicating impaired cellular energy status (Fig. 4I). By contrast, HD MSC-, scr MSC-, and FA-corrected fibroblast-conditioned media did not substantially alter ATP content or the relative ATP/AMP balance compared with untreated HD PB-MNCs.

Together, these data indicate that soluble factors released by FA-deficient stromal cells and FA fibroblasts are sufficient to impair mitochondrial respiration, reduce OxPhos efficiency, and induce energetic stress in otherwise HD PB-MNCs.

### 3.6 FA MSCs- and FA fibroblast-conditioned environments induce oxidative stress but not DNA damage in HD MNCs

Given the close relationship between mitochondrial dysfunction and redox imbalance[41,42], antioxidant responses and oxidative damage markers were analysed in HD PB-MNCs exposed to FA-conditioned media. Exposure to shFA MSC- or FA fibroblast-conditioned media increased antioxidant enzyme activity, including GR and CAT, suggesting activation of a compensatory antioxidant response to the oxidative molecules provided by the medium (Fig. 5A and 5B). Despite this response, HD PB-MNCs exposed to FA-conditioned media accumulated oxidative damage, with increased nitrotyrosine, MDA, and 4-HNE levels (Fig. 5C–E).

**Figure 5.**
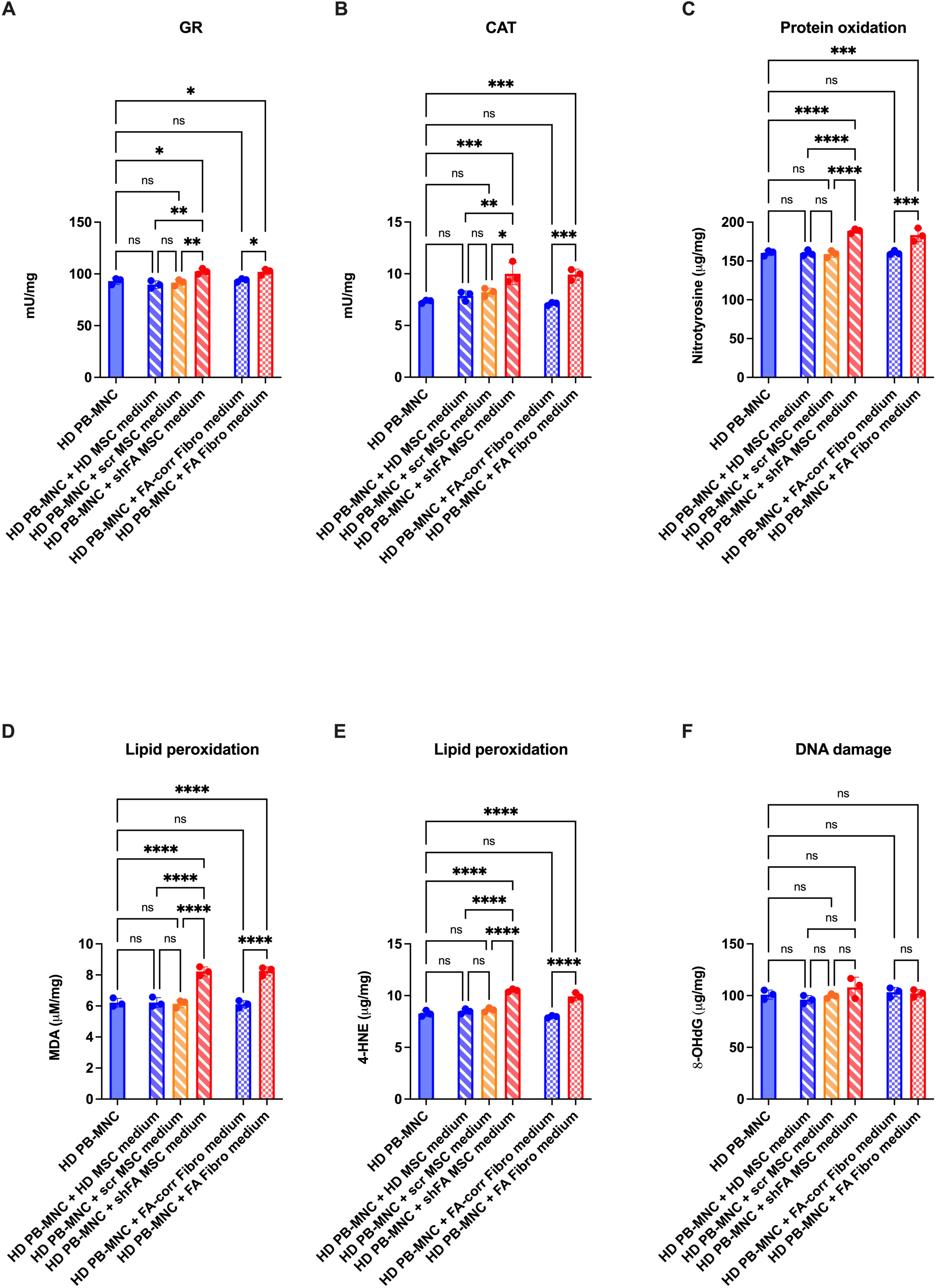
Antioxidant enzyme activity and oxidative damage markers in HD MNCs exposed to stromal-cell-conditioned media. Healthy donor peripheral blood mononuclear cells (HD PB-MNCs) were cultured either untreated or in conditioned media derived from HD mesenchymal stromal cells (HD MSC medium), scramble-control MSCs (scr MSC medium), FANCA-silenced MSCs (shFA MSC medium), FANCA-corrected fibroblasts (FA-corr fibro medium), or FANCA-mutated fibroblasts (FA fibro medium). (A) Glutathione reductase activity. (B) Catalase activity. (C) Protein oxidation, assessed as nitrotyrosine levels. (D) Lipid peroxidation, assessed as malondialdehyde levels. (E) Lipid peroxidation assessed as 4-hydroxynonenal levels. (F) DNA oxidative damage assessed as 8-hydroxy-2ʹ-deoxyguanosine (8OHdG) levels. Data are presented as mean ± SD (n= 3 for each panel). Statistical analysis was performed using one-way ANOVA, followed by Tukey’s post hoc multiple comparison test. ns, not significant; *p < 0.05, **p < 0.01, ***p < 0.001, ****p < 0.0001.

Interestingly, 8-OHdG levels were not significantly altered, indicating that, under these experimental conditions, oxidative damage primarily involved lipids and proteins rather than detectable DNA oxidation (Fig. 5F). Conversely, the incubation with HD MSCs, scr MSCs, and FA-corr fibroblasts did not alter the redox profile of HD PB-MNCs.

Overall, these results suggest that the FA stromal microenvironment can impose mitochondrial and oxidative stress on hematopoietic cells, leading to lipid and protein oxidation while preserving detectable DNA damage.

### 3.7 The peripheral hematopoietic environment of FA patients after HCT remains pro-inflammatory and pro-oxidative

The persistence of inflammatory and oxidative features of the FA hematopoietic environment after HCT was then assessed. Because BM samples could not be collected after HCT due to ethical and clinical constraints, PB plasma has been analysed as a surrogate compartment to evaluate persistent inflammation and oxidative stress in transplanted FA patients.

Compared with HD plasma, post-HCT FA PB plasma showed increased levels of inflammatory cytokines, including TNF-α, TGF-β, IL-6, and IL-8 (Fig. 6A–D) in samples collected a median of 3.78 years after HCT (Table 1), indicating persistence of a pro-inflammatory environment despite hematopoietic replacement. Conversely, an increase was not observed in post-HCT PB plasma from patients with Severe Aplastic Anemia (SAA) that were collected earlier after HCT (1.78 years, Table 1) as compared with FA samples.

**Figure 6.**
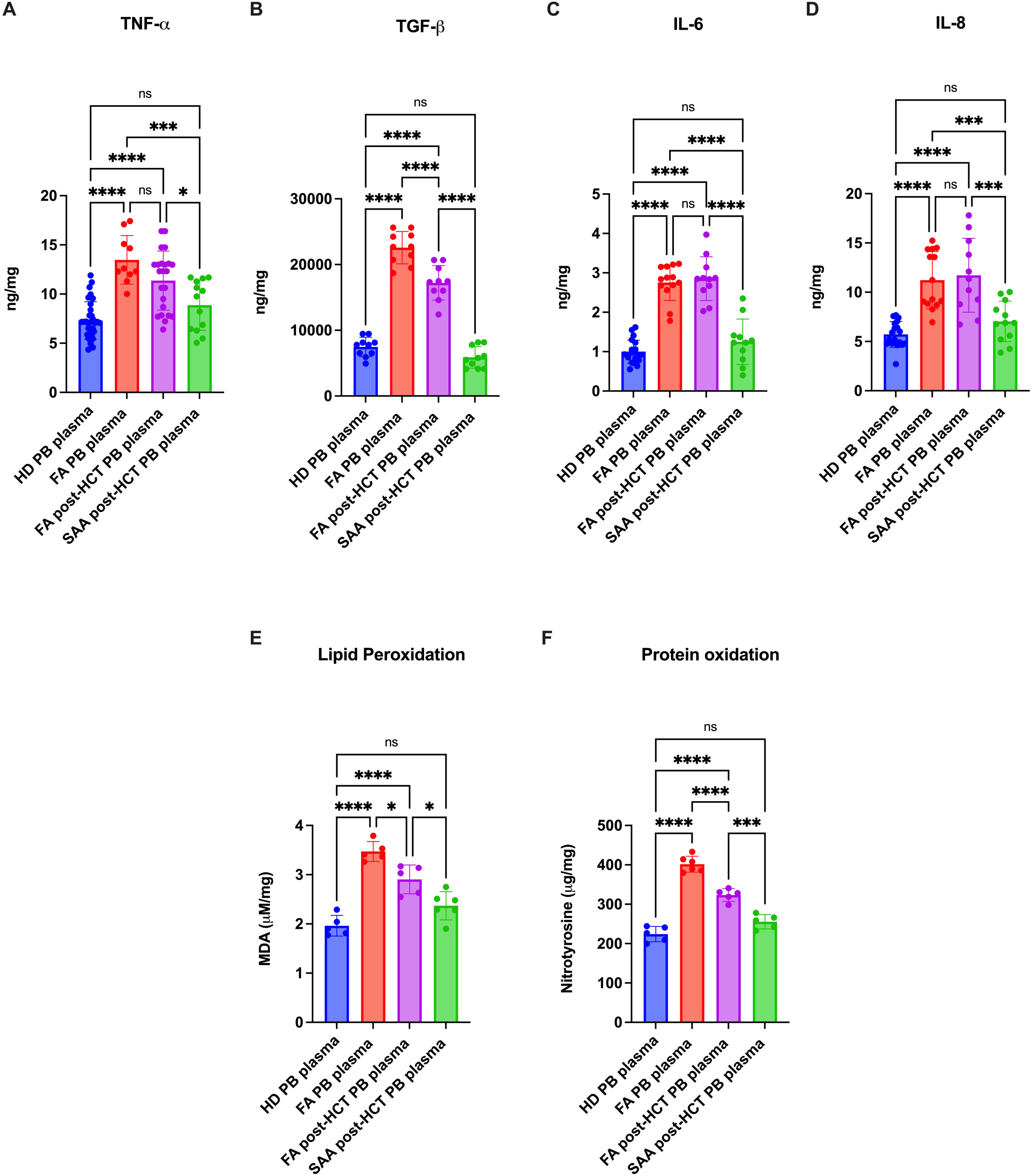
Inflammatory and oxidative damage in PB plasma from FA patients after HCT. Peripheral blood (PB) plasma was analysed from healthy donors (HD), Fanconi Anemia (FA) patients, FA patients after hematopoietic stem cell transplantation (FA post-HCT), and Severe Aplastic Anemia patients after hematopoietic stem cell transplantation (SAA post-HCT). (A–D) Plasma levels of TNF-α, TGF-β, IL-6, and IL-8, respectively. (E) Plasma lipid peroxidation, assessed as malondialdehyde (MDA) levels. (F) Plasma protein oxidation, assessed as nitrotyrosine levels. Data are presented as mean ± SD (n≥8 for A-D panels and n= 5 for E-F panels). Statistical analysis was performed using one-way ANOVA, followed by Tukey’s post hoc multiple comparison test. ns, not significant; *p < 0.05, ***p < 0.001, ****p < 0.0001.

Consistent with the cytokine profile, oxidative stress markers remained elevated in FA PB plasma long after HCT, with increased MDA and nitrotyrosine compared with HD and SAA post-HCT plasma. However, levels were generally lower than those observed in non-transplanted FA plasma (Fig. 6E and 6F). The donor– recipient correspondence for the increased levels of MDA and nitrotyrosine is shown in Supplementary Figure 1.

### 3.8 PB-MNCs from FA patients after HCT show mitochondrial dysfunction and reduced cellular energy status

To assess whether the persistent post-HCT FA environment affects energy metabolism of donor-derived hematopoietic cells, mitochondrial function has been analysed in PB-MNCs from FA post-HCT patients and compared with HD and SAA post-HCT, as controls, and FA subjects.

FA post-HCT PB-MNCs showed a significant reduction in ATP synthesis, OCR, and OxPhos efficiency when pyruvate/malate (P/M) was used as substrate (Fig. 7A–C). However, this impairment was less pronounced than that observed in FA PB-MNCs before transplantation. Interestingly, in FA post-HCT PB-MNCs, metabolic alterations were also evident under succinate-driven respiration, indicating that both complex I- and complex II-linked mitochondrial pathways were affected (Fig. 7D–F). This differed from FA PB-MNCs, in which succinate-linked metabolism was increased compared with controls, suggesting a compensatory response to complex I-linked OxPhos dysfunction. In FA post-HCT PB-MNCs, mitochondrial dysfunction was associated with reduced ATP levels and a decreased ATP/AMP ratio, whereas AMP levels were not consistently increased compared with HD and SAA post-HCT samples (Fig. 7G–I). By contrast, FA PB-MNCs showed increased AMP content, which further contributed to the marked reduction in cellular energy status. The donor–recipient correspondence for the OxPhos function readouts is shown in Supplementary Figure2.

**Figure 7.**
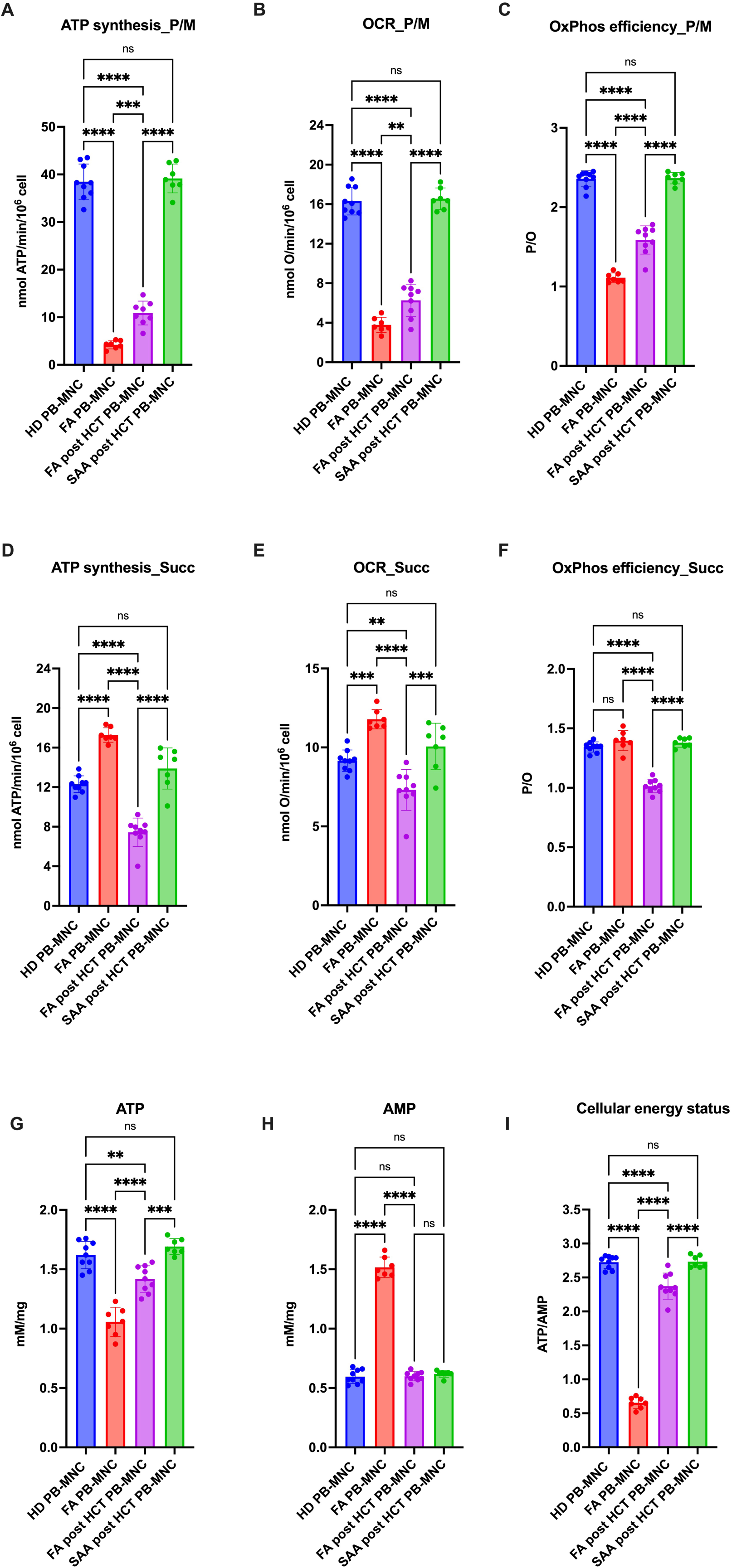
Mitochondrial respiration and cellular energy status in PB-MNCs from FA patients before and after HCT. Peripheral blood mononuclear cells from healthy donors (HD PB-MNC), non-transplanted Fanconi Anemia (FA PB-MNC) patients, FA patients after hematopoietic stem cell transplantation (FA post-HCT PB-MNC), and Severe Aplastic Anemia patients after hematopoietic stem cell transplantation (SAA post-HCT PB-MNC) were analysed for mitochondrial function and energetic status. (A–C) ATP synthesis, oxygen consumption rate (OCR), and OxPhos efficiency, expressed as P/O ratio, were measured under pyruvate/malate-driven (P/M) respiration. (D–F) ATP synthesis, OCR, and OxPhos efficiency, expressed as P/O ratio, were measured under succinate-driven (Succ) respiration. (G) Intracellular ATP levels. (H) Intracellular AMP levels. (I) Cellular energy status expressed as the ATP/AMP ratio. Data are presented as mean ± SD (n≥6 for A-F panels and n = 7 for G-I panels). Statistical analysis was performed using one-way ANOVA, followed by Tukey’s post hoc multiple comparison test. ns, not significant; *p < 0.05, ***p < 0.001, ****p < 0.0001.

Overall, these data indicate that, despite hematopoietic replacement, donor-derived PB-MNCs from FA recipients acquire mitochondrial and energetic defects after transplantation. Importantly, these defects appear mechanistically distinct from those observed in FA PB-MNCs before HCT: in the pre-transplant setting, the defect mainly reflects FANCA-mutated hematopoietic cells with complex I-linked impairment and compensatory succinate-driven respiration, whereas after HCT it involves donor-derived cells exposed to the FA recipient microenvironment and affects both complex I- and complex II-linked respiration.

### 3.9 FA post-HCT PB-MNCs show chronic antioxidant activation with residual oxidative damage

We then investigated whether the metabolic impairment of FA post-HCT PB-MNCs was associated with altered antioxidant response and oxidative damage. Compared with HD and SAA post-HCT PB-MNCs, FA post-HCT cells showed significantly increased glutathione reductase and catalase activities (Fig. 8A and 8B), suggesting activation of an antioxidant response in transplanted FA recipients.

**Figure 8.**
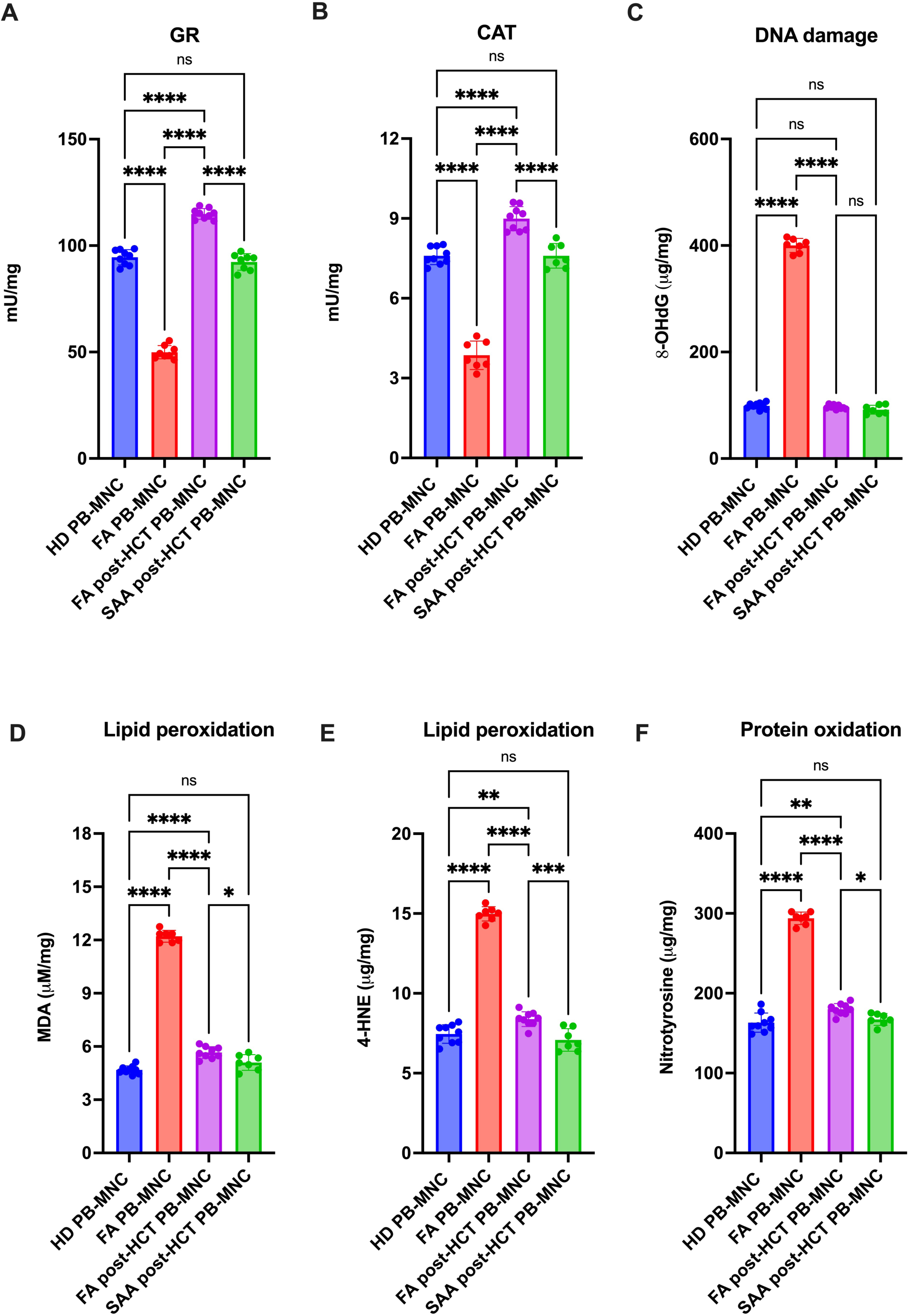
Oxidative damage markers and antioxidant enzyme activities in PB-MNCs from FA patients after HCT. Peripheral blood mononuclear cells from healthy donors (HD PB-MNC), non-transplanted Fanconi Anemia (FA PB-MNC) patients, FA patients after hematopoietic stem cell transplantation (FA post-HCT PB-MNC), and Severe Aplastic Anemia patients after hematopoietic stem cell transplantation (SAA post-HCT PB-MNC) were analysed for antioxidant enzyme activity and oxidative damage markers. (A) Glutathione reductase (GR) activity. (B)Catalase (CAT) activity. (C) DNA oxidative damage assessed as 8-hydroxy-2ʹ-deoxyguanosine (8-OHdG) levels. (D, E) Lipid peroxidation, assessed as malondialdehyde (MDA) and 4-hydroxynonenal (4-HNE) levels, respectively. (F) Protein oxidation, assessed as nitrotyrosine levels. Data are presented as mean ± SD (n = 7 for each panel). Statistical analysis was performed using one-way ANOVA, followed by Tukey’s post hoc multiple comparison test. ns, not significant; *p < 0.05, **p < 0.01, ***p < 0.001, ****p < 0.0001.

Despite this increased antioxidant activity, FA post-HCT PB-MNCs displayed elevated lipid and protein oxidation, as shown by increased MDA, 4-HNE, and nitrotyrosine levels (Fig. 8D–F). In contrast, 8-OHdG levels were not significantly increased compared with HD and SAA post-HCT controls (Fig. 8C). FA PB-MNCs showed a more severe oxidative phenotype, characterized by marked accumulation of DNA, lipid, and protein oxidative damage (Fig. 8). By contrast, they displayed reduced antioxidant activity compared to FA post-HCT PB-MNCs. These findings suggest that after HCT, donor-derived PB-MNCs in FA recipients are chronically exposed to oxidative stress and activate antioxidant defences; however, this response is not sufficient to fully prevent oxidative damage to lipids and proteins. The donor–recipient correspondence for the OxPhos function readouts is shown in Supplementary Figure 3.

Importantly, these metabolic and oxidative alterations remained evident after 96 h of *ex vivo* culture, with FA post-HCT PB-MNCs retaining impaired OxPhos efficiency, reduced cellular energy status, increased antioxidant enzyme activity, and persistent lipid/protein oxidation (Supplementary Fig. 4A–I).

### 3.10 Extracellular vesicles from FA post-HCT patients retain mitochondrial and oxidative alterations

Given that extracellular vesicles (EVs) may reflect and propagate metabolic stress within the circulating microenvironment, mitochondrial function was analysed in PBEVs. EVs from non-transplanted FA patients showed a marked reduction in ATP synthesis, OCR, and OxPhos efficiency compared with HD EVs (Fig. 9A– C). EVs from FA post-HCT patients displayed an intermediate phenotype, with mitochondrial parameters partially reduced compared with those of HD and SAA post-HCT EVs. Still, they were less severely affected than FA EVs (Fig. 9A–C). EVs isolated from FA post-HCT plasma also retained increased lipid peroxidation compared with HD and SAA post-HCT EVs, although again less pronounced than in EVs from non-transplanted FA patients (Fig. 9D).

**Figure 9.**
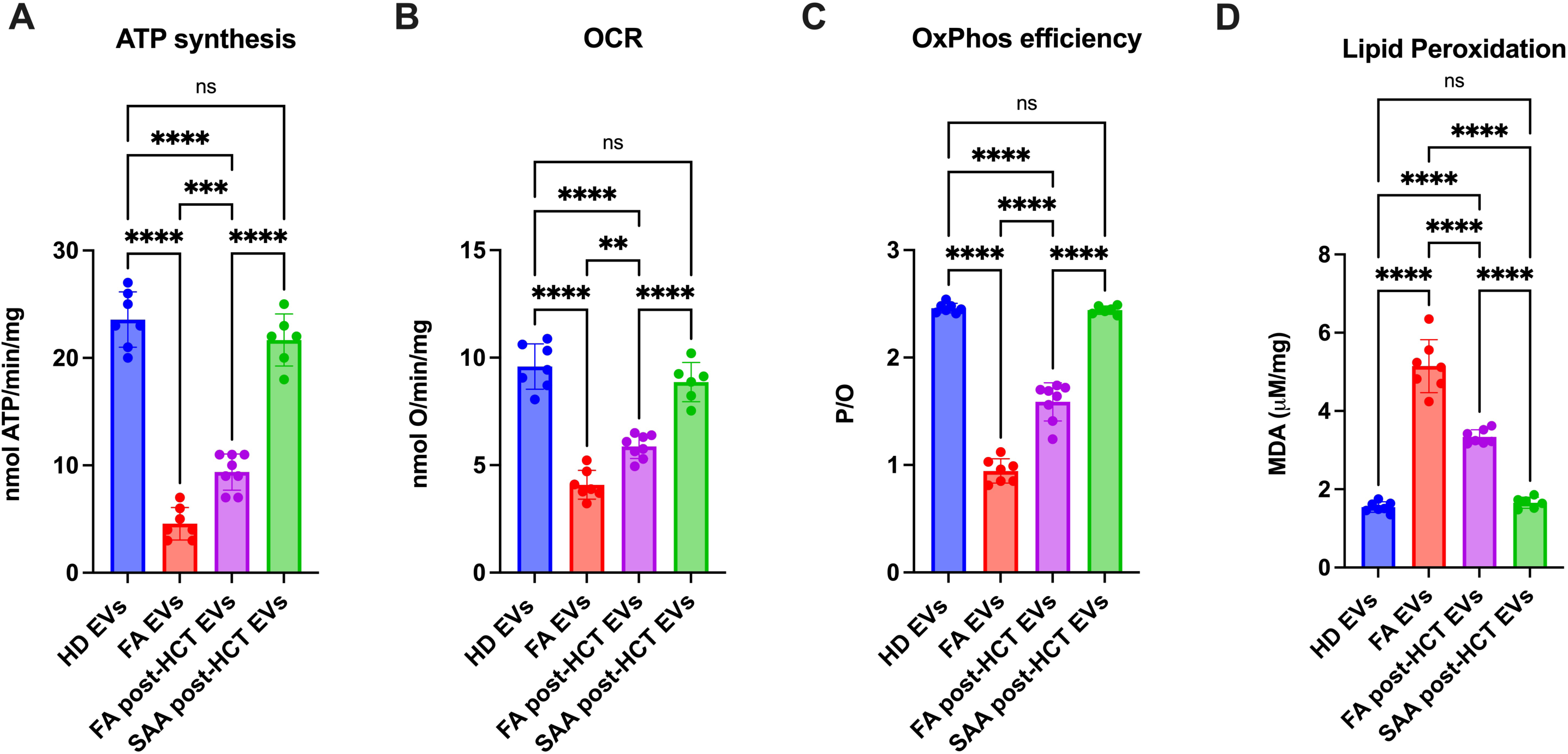
Bioenergetic profiling of extracellular vesicles from FA patients after HCT. Mitochondrial function was assessed in extracellular vesicles (EVs) isolated from healthy donors (HD EVs), non-transplanted Fanconi Anemia patients (FA EVs), Fanconi Anemia patients after hematopoietic stem cell transplantation (FA post-HCT EVs), and Severe Aplastic Anemia patients after hematopoietic stem cell transplantation (SAA post-HCT EVs). (A) ATP synthesis rate, expressed as nmol ATP/min/mg. (B) Oxygen consumption rate (OCR), expressed as nmol O/min/mg. (C) Oxidative phosphorylation efficiency, calculated as the P/O ratio. (D) Lipid peroxidation in PB-derived EVs, assessed as MDA levels. Data are presented as mean ± SD (n≥6 for each panel). Statistical analysis was performed using one-way analysis of variance (ANOVA), followed by Tukey’s post hoc multiple comparison test. ns, not significant; **p < 0.01, ***p < 0.001, ****p < 0.0001.

These data indicate that the extracellular compartment of FA patients is still metabolically altered after HCT, despite hematopoietic replacement, and suggest that EVs may contribute to the maintenance of a dysfunctional circulating microenvironment.

### 3.11 Donor-derived PB-MNCs from FA recipients acquire an altered DNA methylation profile after HSCT

To determine whether the FA recipient environment affects the epigenetic profile of donor-derived hematopoietic cells, DNA methylation profiles in FA patients before and after HSCT and in their healthy donors were investigated. Principal component analysis (PCA) based on genome-wide methylation data (all CpGs included in the methylation array, Figure 10A) revealed a significant separation between pre-HSCT FA samples and donor-derived cells along PC1 (Wilcoxon test, p < 0.0001). In contrast, post-HSCT samples clustered closely with donor samples, with no significant difference observed between donor and post-HSCT profiles (p > 0.05), suggesting that the global DNA methylation landscape of hematopoietic cells is largely restored following transplantation. We then performed a PCA restricted to the CpGs comprising the diagnostic episignature previously described by Pagliara et al.[43] (Figure 10B). In this context, post-HSCT samples displayed an intermediate methylation profile between pre-HSCT FA samples and healthy donors.

**Figure 10.**
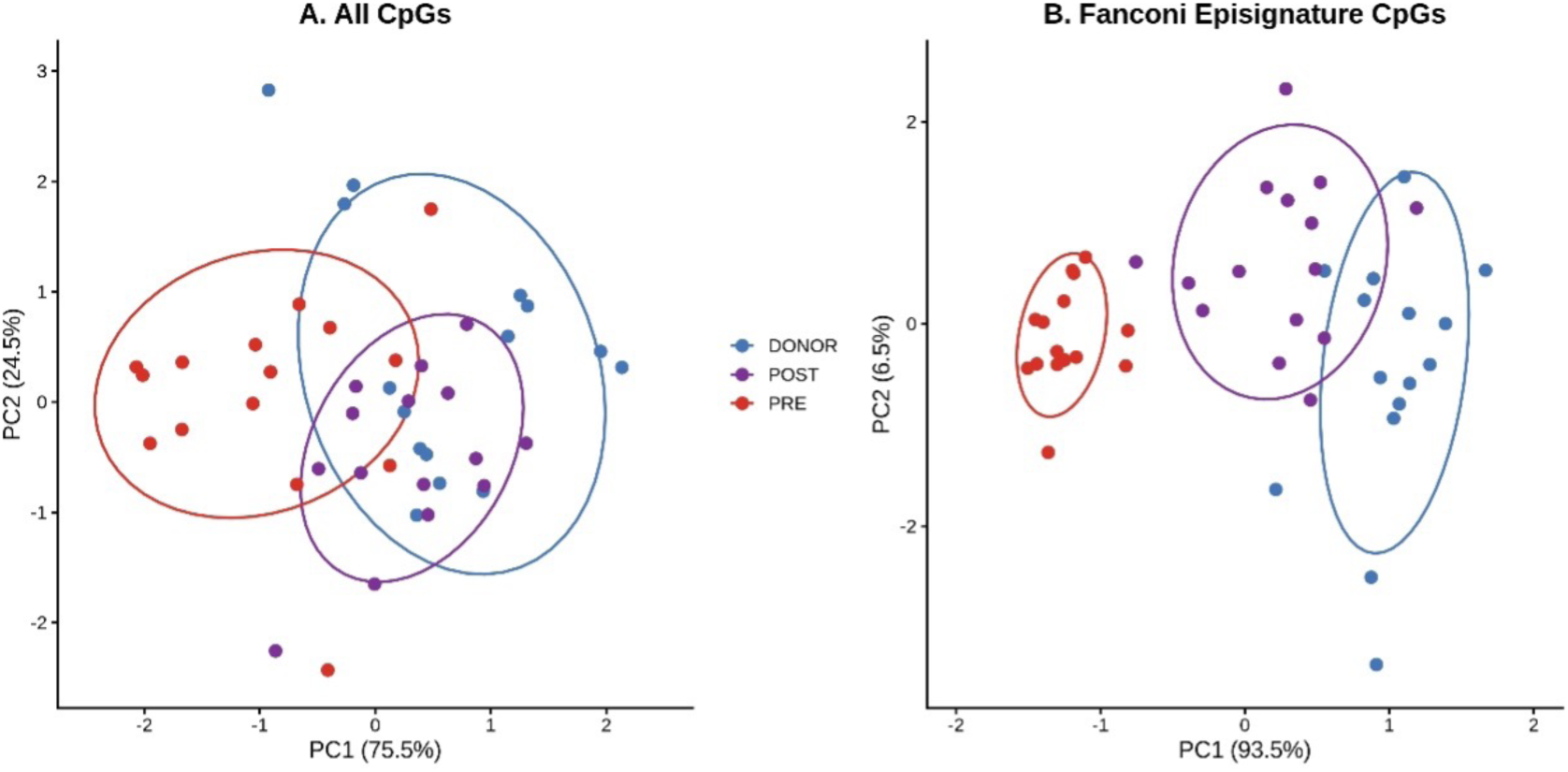
Principal component analysis of DNA methylation profiles in FA samples. PCA was performed using DNA methylation M-values from FA samples collected before hematopoietic stem cell transplantation (PRE, red), after transplantation (POST, purple), and from their corresponding healthy donors (DONOR, blue). (A) PCA based on all CpGs showed substantial overlap among POST and DONOR groups, with a significant difference on the first PC for the PRE samples. (B) PCA restricted to the published Fanconi Anemia episignature. CpGs revealed a clearer separation of the three groups. Ellipses represent the 95% confidence interval for each group.

Pairwise comparisons among the three groups showed statistically significant differences for all comparisons (p < 0.0001), indicating that, although transplantation substantially shifts the methylation profile toward the donor state, the FA-associated episignature is not completely reverted after HSCT and retains features influenced by the recipient environment.

This result indicates that donor-derived PB-MNCs acquire a distinct methylation profile after transplantation into FA recipients. Although this analysis does not establish causality by itself, the intermediate positioning of FA post-HSCT samples is consistent with an epigenetic adaptation of donor hematopoietic cells to the FA recipient microenvironment.

To determine whether the persistent inflammatory phenotype observed after HSCT may be attributable, at least in part, to an epigenetic memory influencing protein regulation, DNA methylation-derived episcores for a subset of circulating proteins selected based on their established roles in inflammation, oxidative stress, stromal remodeling, and endothelial activation were investigated (Table II).

**Table II.**
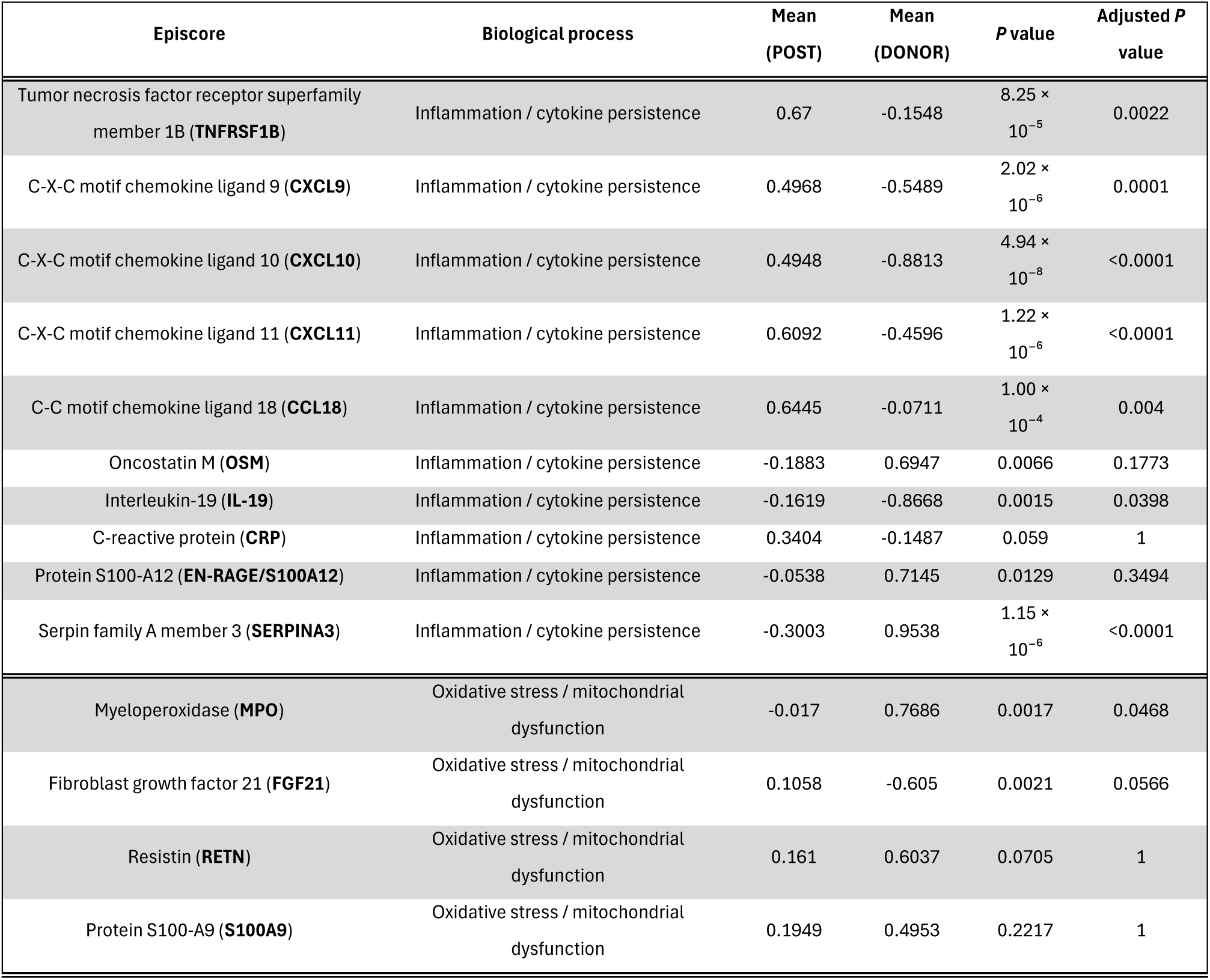
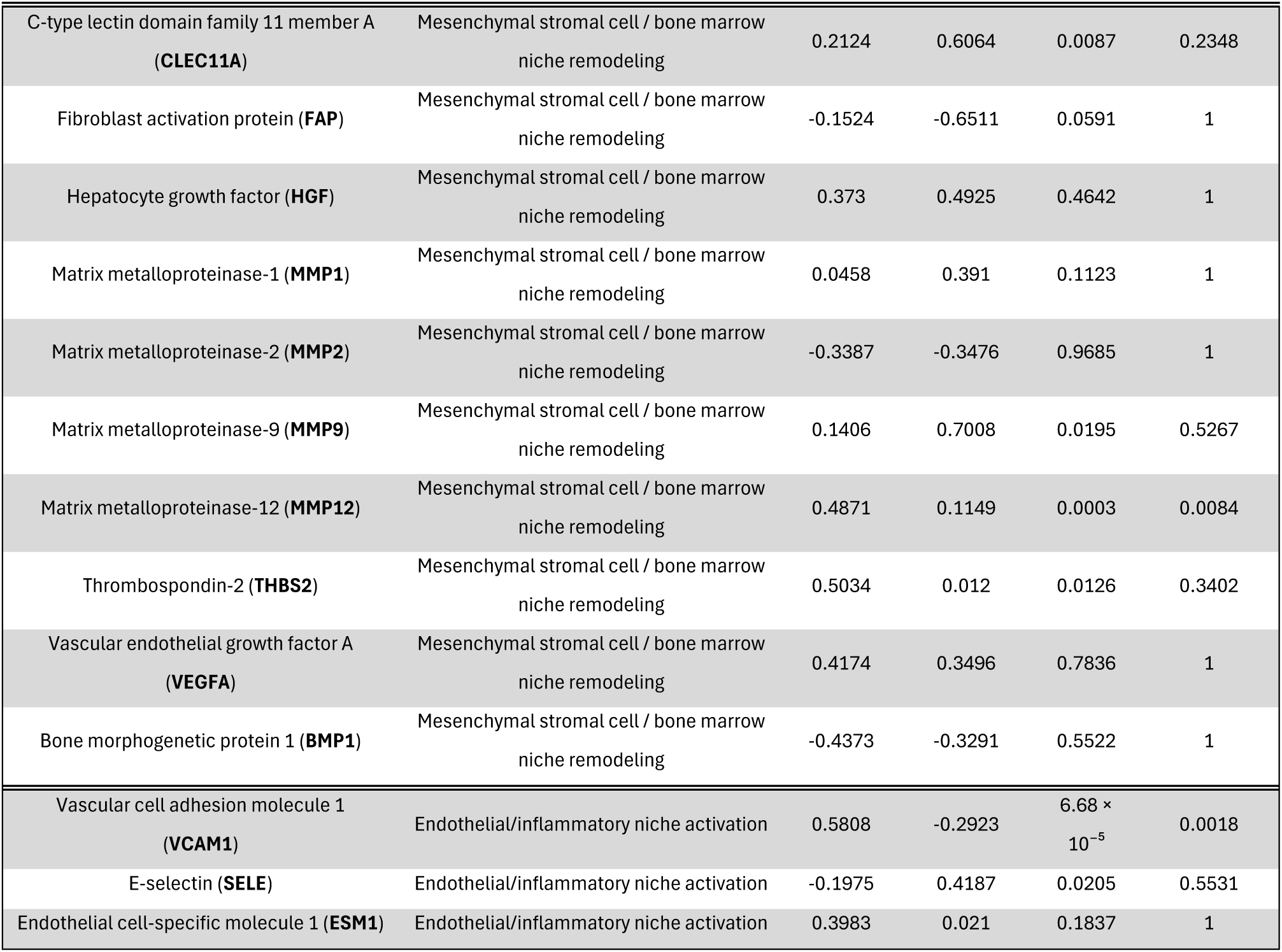
DNA methylation-derived episcores for a subset of circulating proteins selected based on their established roles in inflammation, oxidative stress, stromal remodeling, and endothelial activation.

Compared with paired HD, post-HSCT FA recipients exhibited significant alterations in multiple proteins associated with Inflammation/cytokine persistence, including TNFRSF1B, CXCL9, CXCL10, CXCL11, CCL18, IL-19, and SERPINA3, whereas CRP, OSM, and EN-RAGE did not remain significantly modified after multiple-testing correction. Among proteins related to oxidative stress and mitochondrial dysfunction, only myeloperoxidase (MPO) remained significantly altered after adjustment for multiple comparisons. Markers associated with bone marrow niche remodelling showed more limited changes, with significant differences detected for matrix metalloproteinase-12 (MMP12), while CLEC11A, FGF21, MMP9, and thrombospondin-2 (THBS2) displayed nominal significance but did not withstand multiple-testing correction. Finally, among markers of endothelial and inflammatory niche activation, vascular cell adhesion molecule-1 (VCAM1) remained significantly increased in FA recipients after adjustment, whereas E-selectin and endothelial cell-specific molecule-1 (ESM1) were not significantly different.

Overall, these findings indicate that, despite successful hematopoietic engraftment, FA recipients retain an epigenetic signature predictive of altered regulation of proteins involved predominantly in inflammatory signalling, together with pathways related to oxidative stress, extracellular matrix remodelling, and endothelial activation. These methylation-derived protein episcores support the presence of persistent recipient-driven epigenetic remodelling and are consistent with the pro-inflammatory and pro-oxidative environment identified by the functional and DNA methylation analyses.

### 3.12 FA post-HCT PB-MNCs show chronic activation of stress-response pathways and reduced adaptive capacity to acute stimulation

Based on the evidence of persistent oxidative and inflammatory stress after HCT, the ability of FA post-HCT PB-MNCs to respond to an acute stress stimulus was assessed. Cells were treated with R848, a TLR7/8 agonist used to mimic viral/inflammatory stimulation.

R848 induced glutathione reductase and catalase activity in both HD and FA post-HCT PB-MNCs. However, the relative increase induced by R848 was significantly lower in FA post-HCT cells compared with HD cells, indicating a reduced adaptive antioxidant reserve (Fig. 11A and 11B). In parallel, R848 induced a stronger accumulation of oxidative damage in FA post-HCT PB-MNCs, with increased lipid peroxidation and DNA oxidation compared with stimulated HD cells (Fig. 11C and 11D). R848 also reduced cellular energy status, as shown by a decreased ATP/AMP ratio, with a more pronounced impairment in FA post-HCT PB-MNCs (Fig. 11E).

**Figure 11.**
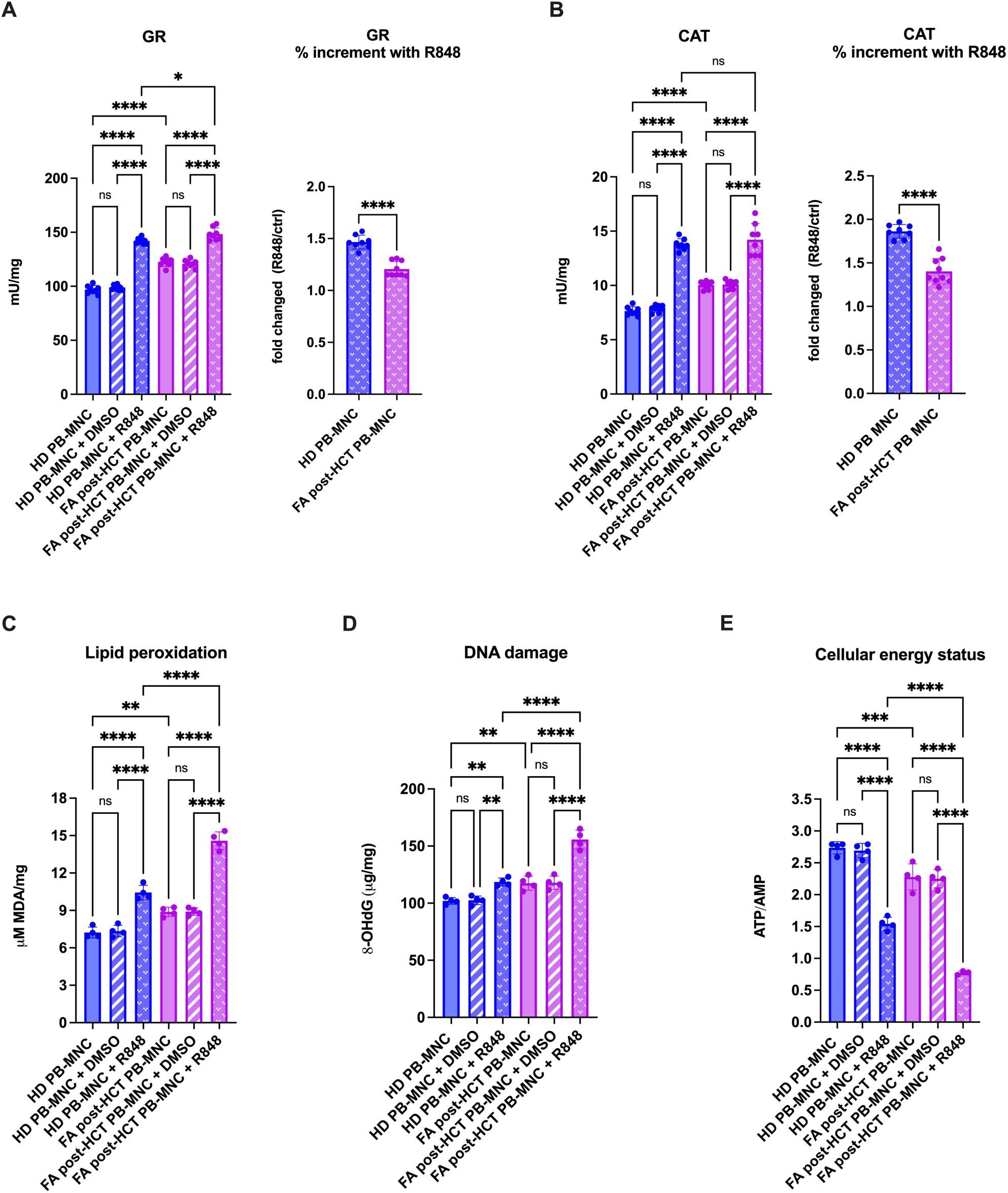
Redox and metabolic response of PB-MNCs to acute R848 stimulation after HCT. Peripheral blood mononuclear cells (PB-MNC) from healthy donors (HD) and Fanconi Anemia patients after hematopoietic cell transplantation (FA post-HCT) were analyzed under basal conditions and following exposure to R848 or to DMSO, used as vehicle control. (A) Glutathione reductase (GR) activity. (B) Catalase (CAT) activity. (C) Lipid peroxidation, measured as MDA levels. (D) Oxidative DNA damage measured as 8-OHdG levels. (E) Cellular energy status, expressed as the ATP/AMP ratio. Data are presented as mean ± SD (n≥4 for each panel). Statistical analysis was performed using one-way analysis of variance (ANOVA), followed by Tukey’s post hoc multiple comparison test for each panel, except for the percentage of increment with R848 for GR and CAT, which was analysed by paired Student’s t-test. ns, not significant; *p < 0.05, **p < 0.01, ****p < 0.0001.

Western blot analysis further supported the presence of chronic stress-response activation in FA post-HCTPB-MNCs. NRF2 expression in peripheral blood (PB) mononuclear cells (MNCs) of FA patients after HCT (FA post-HCT) showed a significant increment compared with HD PB-MNCs, consistent with sustained antioxidant pathway activation (Fig. 12A and 12B). In addition, phosphorylation of STAT2 and SMAD3 was significantly increased in FA post-HCT PB-MNCs, indicating chronic activation of inflammatory signalling pathways associated with interferon-related and TGF-β-related responses (Fig. 12A, 12C, and 12D).

**Figure 12.**
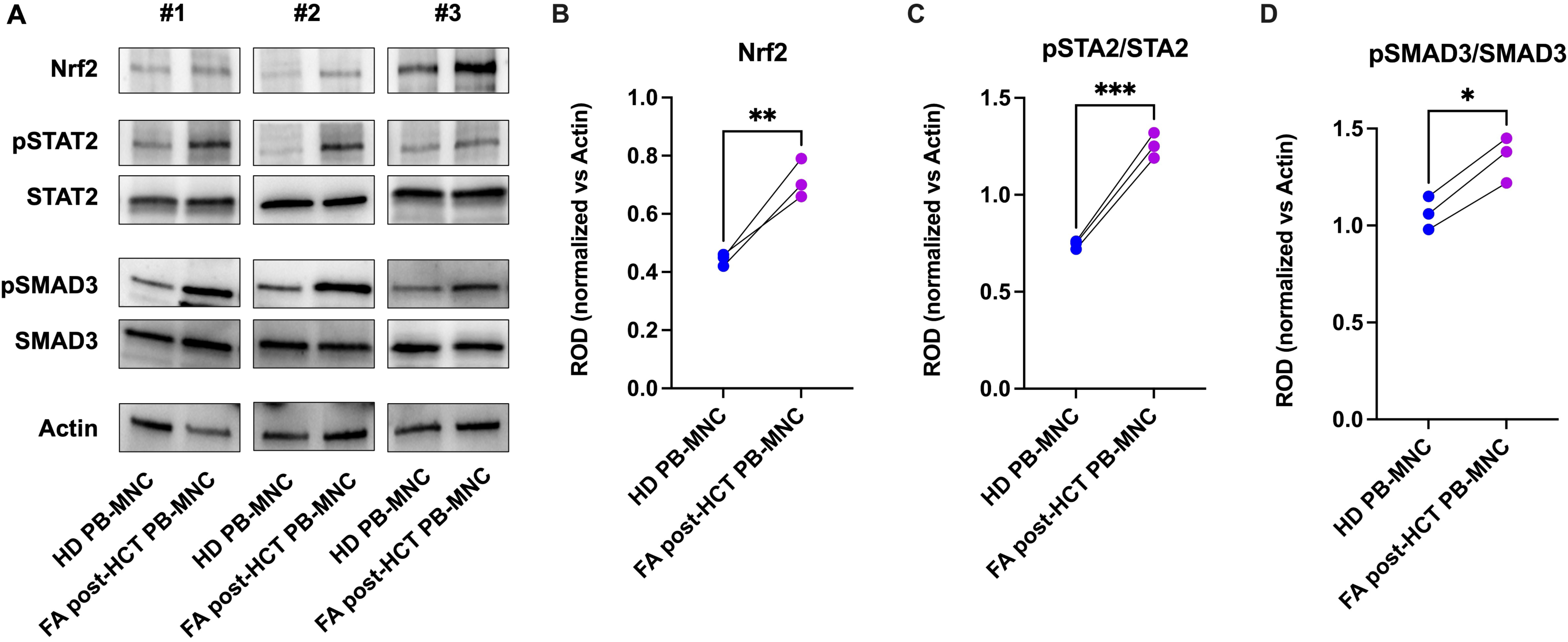
Activation of Nrf2, STAT2, and SMAD3 signalling pathways in PB-MNCs from FA patients after HCT. (A) Representative Western blot analysis of Nrf2, phosphorylated STAT2 (pSTAT2), total STAT2, phosphorylated SMAD3 (pSMAD3), total SMAD3, and actin (used as housekeeping protein) in PB-MNCs from healthy donors (HD PB-MNCs) and FA patients after HCT (FA post-HCT PB-MNCs). Three independent samples are shown. (B–D) Densitometric quantification of Nrf2 protein levels (B), pSTAT2/STAT2 ratio (C), and pSMAD3/SMAD3 ratio (D), expressed as relative optical density (ROD) after normalization against actin signals. Data are presented as mean ± SD (n = 3 for each panel). Statistical analysis was performed using a paired Student’s t-test. ns, not significant; *p < 0.05, **p < 0.01, ***p < 0.001.

Together, these data indicate that PB-MNCs from FA transplanted patients are chronically primed for oxidative and inflammatory stress responses. Still, this chronic activation is associated with a reduced capacity to adapt to acute inflammatory stimulation.

### 3.13 Inhibition of inflammatory pathways and mitochondrial antioxidant treatment improve energy status and reduce oxidative damage in FA post-HCT PB-MNCs

To assess whether inflammatory signalling and mitochondrial oxidative stress contribute to the altered phenotype of FA post-HCT PB-MNCs, cells were treated with inhibitors targeting TNF-α, TGF-β, and NLRP3-related pathways, or with the mitochondria-targeted antioxidant MitoQ.

Whereas inhibition of TNF-α partially improved reduced cellular energy status, inhibition of TGF-β and NLRP3 produced a stronger recovery, restoring energy status to values comparable to or higher than those observed in HD PB-MNCs. MitoQ also improved the energetic profile of FA post-HCT PB-MNCs, indicating that mitochondrial oxidative stress contributes to the impairment of cellular energy homeostasis (Fig. 13A). All treatments significantly reduced MDA accumulation, with TGF-β inhibition, NLRP3 inhibition, and MitoQ showing the most pronounced effects. TNF-α inhibition also decreased lipid peroxidation, although the reduction appeared less marked than that observed with the other interventions (Fig. 13B).

**Figure 13.**
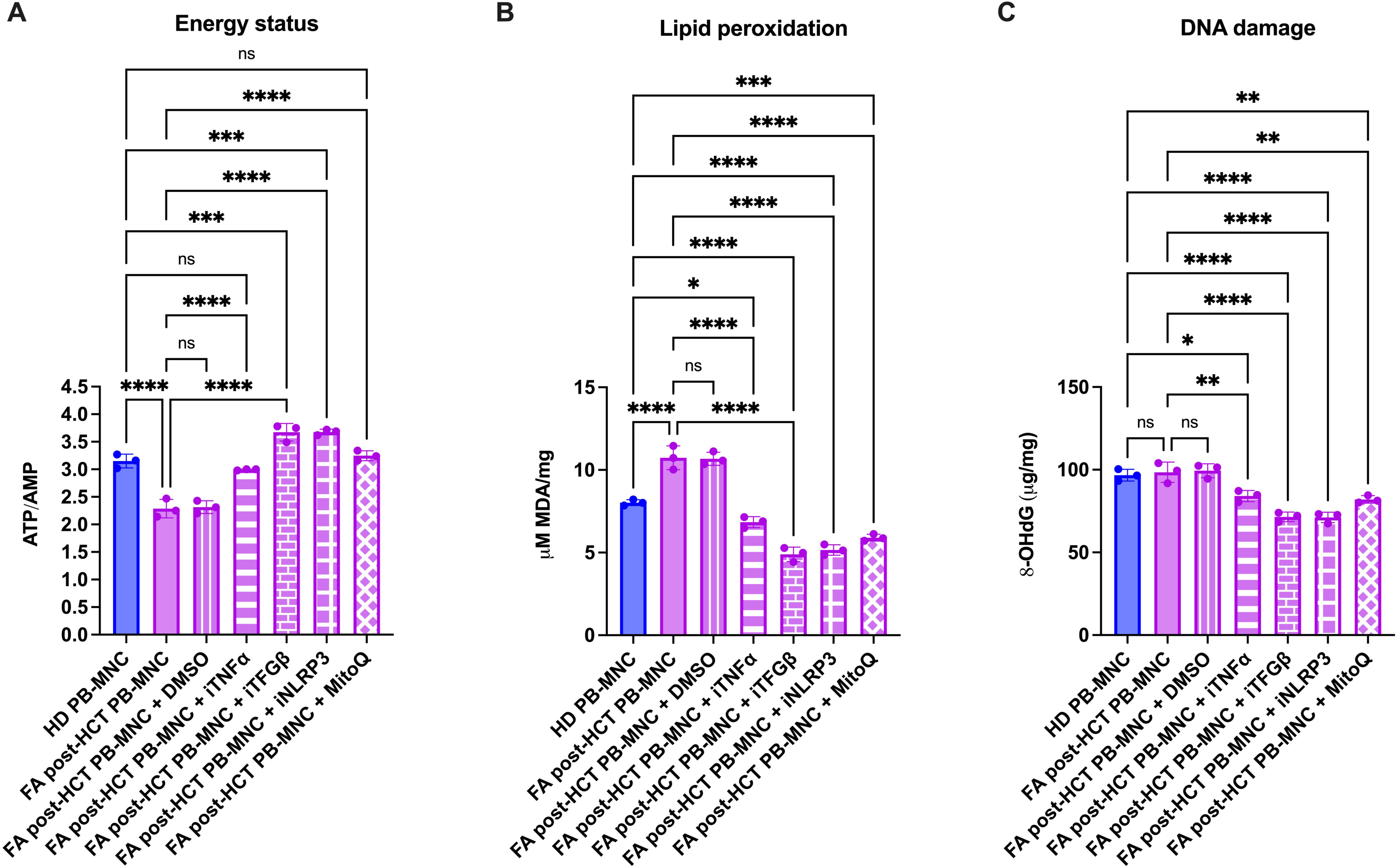
Inhibition of inflammatory pathways and mitochondrial antioxidant treatment modulate energy status and oxidative damage in FA post-HCT PB-MNCs. FA post-HCT PB-MNCs were analysed under basal conditions or after treatment with DMSO, used as vehicle control, or inhibitors targeting TNF-α, TGF-β, and NLRP3-related pathways, or with the mitochondria-targeted antioxidant MitoQ. Results were also compared with HD PB-MNCs. (A) Cellular energy status expressed as ATP/AMP ratio. (B) Lipid peroxidation assessed by MDA levels. (C) Oxidative DNA damage assessed by 8-OHdG levels. Data are presented as mean ± SD (n = 3 for each panel). Statistical analysis was performed using one-way analysis of variance (ANOVA), followed by Tukey’s post hoc multiple comparison test. ns, not significant; *p < 0.05, **p < 0.01, *** p < 0.001, ****p < 0.0001.

Although DNA oxidative damage was not significantly increased in untreated FA post-HCT PB-MNCs compared with HD PB-MNCs, inhibition of inflammatory pathways and MitoQ treatment reduced 8-OHdG levels, suggesting that both inflammatory signalling and mitochondrial oxidative stress contribute to maintaining basal oxidative pressure in donor-derived cells after transplantation (Fig. 13C).

Under R848 stimulation, the same interventions mitigated the exaggerated metabolic and oxidative response of FA post-HCT PB-MNCs. TNF-α inhibition produced a partial rescue, whereas blockade of TGF-β or NLRP3 was more effective, normalizing or increasing the ATP/AMP ratio relative to unstimulated control conditions and markedly limiting lipid peroxidation and oxidative DNA damage. MitoQ also reduced the R848-induced alterations, although its effects on energy status and MDA accumulation were generally weaker than those achieved by TGF-β and NLRP3 inhibition (Supplementary Fig. 2A–C).

Overall, these data indicate that the metabolic and oxidative alterations observed in FA post-HCT PB-MNCs are at least partially reversible. The improvement induced by cytokine-pathway inhibition and by MitoQ supports a model in which chronic inflammatory signalling and mitochondrial oxidative stress cooperate to sustain energetic impairment and oxidative damage in donor-derived mononuclear cells exposed to the FA recipient environment and identifies potential agents able to at least partially restore these altered pathways.

## 4. DISCUSSION

FA is clinically characterized by BM failure caused by the progressive depletion of hematopoietic stem and progenitor cells[1,4,13]. Increasing evidence indicates that this process is not exclusively cell-intrinsic but is also influenced by the BM microenvironment, which regulates HSPC maintenance, proliferation, and differentiation [4,12,28]. In FA, increased levels of cytotoxic and inflammatory cytokines have been detected in both BM and PB plasma, together with hypersensitivity of FA cells to these mediators[5,44]. This inflammatory condition is thought to derive, at least in part, from the DNA repair defect, which promotes genome instability, micronuclei formation, and the release of DNA fragments capable of activating innate inflammatory pathways[10,45]. In parallel, FA cells show high sensitivity to oxidative stress, related to defective mitochondrial metabolism and impaired antioxidant responses[5,8,46]. This defect is particularly relevant during hematopoietic proliferation and differentiation, when the transition from a hypoxic, glycolysis-oriented stem-cell niche to a more oxygen-rich environment requiring increased oxidative phosphorylation may expose FA stem and progenitor cells to severe oxidative stress and apoptosis [6,13,47].

The present study indicates that the hematopoietic environment of FA patients remains persistently pro-oxidative and pro-inflammatory after HCT. This is consistent with the broader view that FA pathophysiology extends beyond the canonical defect in DNA interstrand-crosslink repair and includes chronic oxidative stress, mitochondrial dysfunction, and dysregulated inflammatory signalling, which together may sustain a self-amplifying cycle of cellular damage [8,12,13,48]. A major finding of our work is that donor-derived PB-MNCs from transplanted FA patients do not fully acquire a healthy donor-like phenotype. Instead, they display mitochondrial dysfunction, reduced energetic efficiency, chronic activation of antioxidant and inflammatory pathways, residual oxidative damage, and altered DNA methylation. These observations support the concept that, after HCT, donor hematopoietic cells function within a recipient microenvironment that remains biologically altered. This interpretation is also consistent with recent reviews emphasizing that HCT corrects BM failure but does not eliminate the underlying pro-neoplastic and pro-inflammatory predisposition of FA patients [12].

Inflammation and oxidative stress are closely interrelated and mutually reinforcing processes[41,42]. While cytokine alterations in FA have been widely investigated, extracellular oxidative stress has received less attention. Our findings show increased MDA and nitrotyrosine in BM and PB plasma from FA patients, indicating extracellular lipid and protein oxidation and supporting the presence of a systemic pro-oxidant state extending beyond the hematopoietic compartment [13]. Importantly, MDA and nitrotyrosine should not be interpreted merely as passive markers. Lipid peroxidation products can form protein adducts, alter membrane properties, induce pro-inflammatory cytokines, DNA adducts, and promote immune-cell activation, including T-cell activation [49–51], while nitrotyrosine can modify protein function, exert cytotoxic effects, inhibit cell growth, and disrupt cellular homeostasis [52]. This mechanism is further supported by evidence that the FA pathway participates in antioxidant defence by protecting promoters of major antioxidant genes from oxidative damage, so that its loss directly impairs the cellular antioxidant machinery [53].

Our data further suggest that stromal components of the FA microenvironment actively contribute to this oxidative state. FA MSCs showed impaired mitochondrial metabolism, reduced OxPhos, defective antioxidant enzyme activity, and increased lipid peroxidation, supporting the view that FA stromal cells can release oxidizing activity into the extracellular environment and promote oxidative modification of complex plasma components. This is particularly relevant because MSCs are heterogeneous multipotent stromal cells that regulate BM niche homeostasis and support the maintenance, expansion, and differentiation of hematopoietic stem and progenitor cells [3,54].

BM mesenchymal stromal cells are not eradicated by non-myeloablative conditioning and can persist as host-derived, recipient-origin cells for years after engraftment, as shown by chimerism analyses demonstrating fully recipient-derived MSCs in transplanted FA patients[3]. Thus, HCT replaces the genetically defective hematopoietic compartment but leaves FA-mutant stromal cells and other non-hematopoietic recipient tissues in place. This provides a structural basis for the persistence of a pro-oxidative and pro-inflammatory niche into which donor-derived hematopoietic cells must engraft. In this context, comparison with SAA post-HCT controls is particularly informative, because it suggests that persistent cytokine, oxidative, and mitochondrial abnormalities after transplantation are not simply a generic consequence of HCT but are more closely associated with the FA recipient background.

The functional consequences of this altered microenvironment were demonstrated by exposing HD MNCs to media conditioned by shFA MSCs or FA fibroblasts. These conditions reduced OxPhos efficiency, impaired cellular energy status, and induced oxidative damage in HD cells, indicating that soluble or extracellular factors released by FA stromal cells are sufficient to transfer metabolic and redox stress to otherwise normal hematopoietic cells. This experimental observation provides a mechanistic bridge between the intrinsic defect of FA stromal cells and the altered phenotype observed in donor-derived cells after transplantation. It is also conceptually consistent with previous work showing that the BM niche environment, including low oxygen tension and paracrine signals, strongly modulates the expression of the FA mitochondrial and metabolic phenotype, with pronounced OxPhos and redox abnormalities emerging when cells leave the hypoxic BM niche and enter the oxygen-rich bloodstream environment [6].

Given the close relationship between altered redox homeostasis and inflammatory activation, we analyzed the cytokine profile of PB plasma from HD, FA patients before and after HCT, and SAA patients after HCT. As expected, PB plasma from FA patients showed increased levels of TNF-α, TGF-β, IL-6, and IL-8, as already reported in the literature [12,55–57].

Notably, these cytokines remained elevated in FA post-HCT plasma, whereas comparable increases were not observed in SAA post-HCT samples. This indicates that the phenotype is not merely a generic consequence of transplantation but is associated with the FA recipient background. The persistence of increased cytokine levels long-term after HCT suggests that donor-derived hematopoietic cells remain exposed to inflammatory cues compatible with the metabolic, redox, and signalling abnormalities observed in subsequent experiments. The increased IL-8 levels observed in transplanted FA patients deserve particular attention. IL-8 is involved in inflammation, neutrophil recruitment, angiogenesis, migration, invasion, and tumor-associated processes, and elevated circulating or local IL-8 has been associated with poor prognosis and tumor progression in squamous cell carcinomas, including esophageal and head-and-neck SCC [58]. In the context of FA, where patients are predisposed to squamous cell carcinomas and where this risk persists after otherwise successful HCT [12,59], persistent IL-8 elevation may be biologically relevant. Its increase may reflect, at least in part, chronic oxidative-stress responses and NRF2-related transcriptional activation[60]. However, a direct causal link between NRF2 activation, IL-8 elevation, oxidative stress, and cancer risk cannot be established from the present data and should be addressed in dedicated clinical and mechanistic studies.

In line with the persistence of an inflammatory and oxidative circulating environment, donor-derived PB-MNCs from transplanted FA patients displayed mitochondrial dysfunction and reduced cellular energy status. Notably, although FA post-HCT PB-MNCs displayed defects in mitochondrial metabolism, these alterations differed from those observed in FA PB-MNCs before transplantation. In FA PB-MNCs, mitochondrial dysfunction was characterized by a severe metabolic impairment, with markedly reduced ATP synthesis, OCR, and P/O ratio under P/M-driven respiration, whereas Succ-driven OxPhos was increased, possibly reflecting a compensatory response. Conversely, SAA post-HCT PB-MNCs were largely comparable to HD controls, further supporting the specificity of the metabolic alterations observed in FA post-HCT cells. Therefore, it is possible to speculate that FA post-HCT PB-MNCs showed a distinct metabolic phenotype compared with FA PB-MNCs, with less pronounced alterations in some parameters but still significant impairment compared with HD and SAA post-HCT controls. In other words, this suggests that mitochondrial dysfunction in FA post-HCT PB-MNCs reflects additional mechanisms related to the FA recipient environment. The reduction in P/O ratio and ATP/AMP ratio further indicates energetic stress and less efficient oxidative phosphorylation after engraftment in FA recipients.

Despite increased antioxidant enzyme activity, likely reflecting an adaptive response to the inflammatory and oxidative milieu, FA post-HCT PB-MNCs still accumulated oxidative damage. This residual damage may derive both from persistent extracellular oxidative stress and from altered mitochondrial metabolism, potentially establishing a self-sustaining cycle in which mitochondrial dysfunction and oxidative damage reinforce each other. Importantly, these abnormalities were less severe than those observed in non-transplanted FA cells but remained evident compared with healthy donor and SAA post-HCT controls. This intermediate phenotype is consistent with the interpretation that donor-derived cells are genetically normal but are functionally remodelled by the FA recipient environment.

Notably, these metabolic and oxidative alterations remained evident after 96 h of ex vivo culture, suggesting that exposure to the FA recipient microenvironment induces a relatively stable metabolic and redox imprinting in donor-derived PB-MNCs that is not rapidly reversed after removal from the in vivo environment. The methylation data provide a further layer of interpretation to this persistent phenotype. Oxidative stress, inflammation, mitochondrial metabolism, and epigenetic regulation are closely interconnected: chronic inflammation can promote oxidative damage, transcriptional dysregulation, and cellular dysfunction, whereas mitochondrial function provides intermediate metabolites required for the generation and modification of nuclear epigenetic marks through mitochondria–nucleus communication. Consistent with this concept, donor-derived cells from FA recipients did not cluster completely with donor samples but occupied an intermediate position between donor and FA pre-HCT profiles. Although these data do not prove causality, they support the hypothesis that the FA recipient environment contributes to epigenetic remodelling of donor cells. Similar findings have been reported in other allogeneic HCT settings, where donor hematopoietic stem/progenitor cells did not fully retain their pre-transplant methylation profile after engraftment, but acquired a transient, partially reversible hypermethylation signature interpreted as donor-cell adaptation to the recipient bone marrow niche, affecting genes involved in hematopoietic activation, differentiation, and trafficking[61]. Therefore, the altered methylation profile observed in our FA cohort may represent a disease-specific instance of recipient-driven epigenetic remodelling of donor-derived hematopoietic cells under chronic microenvironmental stress.

The chronic stress phenotype was further supported by increased NRF2 expression in FA post-HCT PB-MNCs, consistent with compensatory activation of antioxidant responses following FA pathway dysfunction[53]. Increased STAT2 and SMAD3 phosphorylation also indicated persistent interferon-and TGF-β-related signalling, in agreement with evidence linking FA genome instability to cGAS/STING–type-I interferon activation and premature inflammatory aging [10,62] and showing that hyperactive TGF-β/SMAD3 signalling impairs DNA repair and hematopoietic function in FA[4]. However, chronic activation of antioxidant and inflammatory pathways does not necessarily imply improved stress resistance. When challenged with R848, used to mimic an acute inflammatory or infection-like stimulus, FA post-HCT PB-MNCs showed a lower relative increase in antioxidant enzyme activity and greater accumulation of oxidative damage than donor cells. This suggests a reduced adaptive reserve, possibly because antioxidant and inflammatory pathways are already chronically engaged under basal conditions. Thus, donor-derived cells from FA recipients appear to be persistently activated, but less capable of mounting an efficient additional response to acute stress.

The rescue experiments with cytokine inhibitors and antioxidant treatment further support the functional relevance of inflammatory and oxidative pathways. Targeting TNF-α, TGF-β, or NLRP3-related inflammatory signalling improved energy status and reduced oxidative damage in FA post-HCT PB-MNCs. These findings suggest that the mitochondrial and redox abnormalities observed in donor-derived cells are not irreversible and may be at least partly driven by modifiable extracellular signals. This is consistent with the established role of TNF-α overproduction in suppressing FA hematopoiesis [55] and with the contribution of hyperactive TGF-β/SMAD3 signalling to FA-associated hematopoietic stem cell dysfunction [4]. Nevertheless, these data should be interpreted as mechanistic support rather than definitive therapeutic proof. Further studies will be required to define the dominant upstream mediators, the relative contribution of each pathway, and whether similar effects can be achieved *in vivo*.

Two major limitations should be acknowledged. First, post-HCT analyses were performed on peripheral blood plasma and PB-MNCs because bone marrow aspirates after transplantation were not available for ethical and clinical reasons. Peripheral blood was therefore used as a surrogate of the hematopoietic environment, but it may not fully recapitulate the bone marrow niche. This caveat is relevant because the metabolic and redox phenotype of FA cells is strongly influenced by oxygen tension and may be partly masked in the hypoxic bone marrow compartment compared with peripheral blood [6]. Second, the study includes a limited cohort, as expected for a rare disease such as FA, and the results should be validated in larger and longitudinal series. Notably, even the largest published FA-MSC series to date comprised only 28 patients, underscoring that this constraint is structural to the field rather than specific to our study [3]. However, despite these limitations, our study provides evidence that HCT corrects the hematopoietic genetic defect but does not fully normalize the FA recipient microenvironment. Donor-derived hematopoietic cells engraft into a persistently pro-inflammatory and pro-oxidative environment and acquire metabolic, redox, signalling, and epigenetic adaptations. This model may help explain why transplanted FA patients can display persistent biological vulnerability despite hematological correction and is consistent with the clinical observation that HCT does not eliminate the increased cancer risk that characterizes FA [12,59]. It also suggests that long-term management of FA after HCT may require attention not only to donor hematopoiesis, but also to the recipient microenvironment, including the stromal compartment, which remains host-derived and is therefore not corrected by transplantation itself [3]. This may suggest including redox and inflammation status monitoring in post-HCT surveillance plans.

## 5. CONCLUSION

Our findings indicate that the FA hematopoietic microenvironment remains pro-oxidative and pro-inflammatory after HCT and can influence the biology of donor-derived mononuclear cells. Chronic exposure to oxidative and inflammatory stress is associated with mitochondrial dysfunction, altered energy metabolism, persistent NRF2, STAT2, and SMAD3 activation, changes in DNA methylation profile, increased IL-8 levels, and reduced adaptive capacity to acute inflammatory stimulation. These data support a model in which the post-transplant FA recipient environment contributes to long-term remodelling of donor cells, in line with the broader characterization of FA as a disease involving intertwined DNA-repair failure, oxidative stress, mitochondrial dysfunction, and chronic inflammatory signalling[12,13]. Maintaining the balance between oxidative and inflammatory responses may therefore be clinically relevant for preserving cellular homeostasis, including epigenetic control, and oxidative and inflammatory pathways emerge as relevant targets for further mechanistic and translational investigation.

## Supporting information

Whole WB signals

## 6. LIST OF ABBREVIATIONS

4-HNE: 4-hydroxy-2-nonenal
8-OHdG: 8-hydroxy-2ʹ-deoxyguanosine
AMP: adenosine monophosphate
ATP: adenosine triphosphate
BM: bone marrow
CAT: catalase
cGAS: cyclic GMP–AMP synthase
CpG: cytosine–phosphate–guanine
DMSO: dimethyl sulfoxide
EV: extracellular vesicle
FA: Fanconi anaemia
FANCA: Fanconi anaemia complementation group A
GR: glutathione reductase
HCT: haematopoietic cell transplantation
HD: healthy donor
HSC: haematopoietic stem cell
HSPC: haematopoietic stem and progenitor cell
IFN-γ: interferon gamma
MDA: malondialdehyde
MNC: mononuclear cell
MSC: mesenchymal stromal cell
MSD: matched sibling donor
MUD: matched unrelated donor
NLRP3: NLR family pyrin domain containing 3
NRF2: nuclear factor erythroid 2-related factor 2
OCR: oxygen consumption rate
OxPhos: oxidative phosphorylation
P/M: pyruvate/malate
P/O: phosphorylation-to-oxygen ratio
PB: peripheral blood
PCA: principal component analysis
PTCy: post-transplant cyclophosphamide
SAA: severe aplastic anaemia
STAT2: signal transducer and activator of transcription 2
STING: stimulator of interferon genes
TBARS: thiobarbituric acid-reactive substances
TCD: T-cell depletion
TGF-β: transforming growth factor beta
TLR7/8: Toll-like receptor 7/8
TNF-α: tumor necrosis factor alpha

## 7. DECLARATIONS

### Ethics approval and consent to participate

This study was performed in accordance with the Declaration of Helsinki and with the relevant laws, regulatory frameworks, and guidelines applicable at the study site. The study was approved by the Giannina Gaslini Institute Ethics Committee (Approval No. JS 002), initially on 24 September 2010 and most recently following amendments on 21 April 2026. Written informed consent to participate in the study and for publication was obtained from all participants or their legal representatives, as applicable, and participants’ privacy rights were respected. The study included human biological material obtained from donors, for which consent for use in this research and for publication was obtained from the donors or their legal representatives, as applicable.

### Consent for publication

Not applicable

### Availability of data and materials

Data will be made available on request.

### Authors’ contributions

Matilde Balbi: Investigation, Data curation, Formal analysis. Stefano Regis: Investigation, Formal analysis. Giovanni Fiorito: Investigation, Data curation, Methodology, Formal analysis. Paolo Uva: Supervision. Marta Maragliano: Supervision. Fabio Corsolini: Investigation. Federica Raggi: Investigation. Luca Arcuri: Resources. Maria Carla Giarratana: Resources. Sara Pestarino: Resources. Ramona Tallone: Resources.

Andrea Beccaria: Resources.Vanessa Cossu: Investigation. Giorgia Damonte: Investigation. Martina Serra: Investigation. Grazia Maria Spaggiari: Investigation. Carlo Dufour: Funding acquisition, Resources, Writing – review & editing, Supervision. Maura Faraci: Resources. Silvia Ravera: Conceptualization, Data curation, Formal analysis, Funding acquisition, Methodology, Visualization, Resources, Data curation, Writing – original draft, Writing – review & editing, Supervision. Filomena Pierri: Conceptualization, Resources, Writing – review & editing, Supervision. Enrico Cappelli: Conceptualization, Resources, Data curation, Writing – original draft, Writing – review & editing, Supervision. All authors contributed to the interpretation of the data and preparation of the manuscript. All authors read and approved the final version of the manuscript.

## Acknowledgements

We gratefully acknowledge ERG S.p.A., Cambiaso Risso Group, Rimorchiatori Riuniti, Saar Depositi Oleari Portuali, and the Nicola Ferrari ONLUS Foundation for supporting the activities of the Clinical and Experimental Hematology Unit at IRCCS Istituto Giannina Gaslini. We also thank the Gaslini Biobank, IRCCS Istituto Giannina Gaslini, Genoa, a member of the Telethon Network of Genetic Biobanks, for providing the biological specimens used in this study.

## Use of generative AI

During the preparation of this work, the author(s) used ChatGPT for language editing and to improve the clarity and readability of the manuscript. The author(s) reviewed and edited the output as needed and take full responsibility for the content of the published article.

## REFERENCES

1. Ceccaldi R, Parmar K, Mouly E, Delord M, Kim JM, Regairaz M, et al. Bone marrow failure in Fanconi anemia is triggered by an exacerbated p53/p21 DNA damage response that impairs hematopoietic stem and progenitor cells. Cell Stem Cell [Internet]. Cell Stem Cell; 2012 [cited 2024 May 20];11:36–49. 10.1016/J.STEM.2012.05.013

2. Niraj J, Färkkilä A, D’Andrea AD. The fanconi anemia pathway in cancer. Annu Rev Cancer Biol. Annual Reviews Inc.; 2019;3:457–78. 10.1146/ANNUREV-CANCERBIO-030617-050422

3. Zubicaray J, Ivanova M, Iriondo J, Martínez JG, Muñoz-Viana R, Abad L, et al. Role of the mesenchymal stromal cells in bone marrow failure of Fanconi Anemia patients. Front Cell Dev Biol. Frontiers Media SA; 2024;12. 10.3389/FCELL.2024.1286815

4. Zhang H, Kozono DE, O’Connor KW, Vidal-Cardenas S, Rousseau A, Hamilton A, et al. TGF-β Inhibition Rescues Hematopoietic Stem Cell Defects and Bone Marrow Failure in Fanconi Anemia. Cell Stem Cell [Internet]. Cell Stem Cell; 2016;18:668–81. 10.1016/J.STEM.2016.03.002

5. Bagby G. Recent advances in understanding hematopoiesis in Fanconi Anemia. F1000Res. Faculty of 1000 Ltd; 2018;7. 10.12688/F1000RESEARCH.13213.1

6. Cappelli E, Degan P, Bruno S, Pierri F, Miano M, Raggi F, et al. The passage from bone marrow niche to bloodstream triggers the metabolic impairment in Fanconi Anemia mononuclear cells. Redox Biol. 2020;36. 10.1016/j.redox.2020.101618

7. Bertola N, Regis S, Cossu V, Balbi M, Serra M, Corsolini F, et al. miR-29a-3p and TGF-β Axis in Fanconi anemia: mechanisms driving metabolic dysfunction and genome stability. Cell Mol Life Sci [Internet]. Cell Mol Life Sci; 2025;82. 10.1007/S00018-025-05775-W

8. Damonte G, Balbi M, Amaroli A, Cossu V, Panfoli I, Cappelli E, et al. Fanconi Anemia: Interplay Between DNA Repair Defects, Mitochondrial Dysfunction, and Oxidative Stress. Cells. Multidisciplinary Digital Publishing Institute (MDPI); 2026;15. 10.3390/CELLS15090753

9. Saadatzadeh MR, Bijangi-Vishehsaraei K, Hong P, Bergmann H, Haneline LS. Oxidant hypersensitivity of Fanconi anemia type C-deficient cells is dependent on a redox-regulated apoptotic pathway. J Biol Chem [Internet]. J Biol Chem; 2004;279:16805–12. 10.1074/JBC.M313721200

10. Landelouci K, Sinha S, Pépin G. Type-I Interferon Signaling in Fanconi Anemia. Front Cell Infect Microbiol [Internet]. Frontiers Media S.A.; 2022 [cited 2024 Jul 24];12:820273. 10.3389/FCIMB.2022.820273/BIBTEX

11. Nagarajan AG, Neeley N, Doroodian P, Olazabel A, Kupfer G. Genome Wide siRNA Screen Identifies CGAS-Sting Pathway As a Pharmacological Target That Promotes Survival of Hematopoietic Stem Cells Deficient in Fanconi Genes. Blood. American Society of Hematology; 2023;142:1083. 10.1182/BLOOD-2023-187180

12. Repczynska A, Ciastek B, Haus O. New Insights into the Fanconi Anemia Pathogenesis: A Crosstalk Between Inflammation and Oxidative Stress. Int J Mol Sci. Multidisciplinary Digital Publishing Institute (MDPI); 2024;25. 10.3390/IJMS252111619

13. Brosh RM, Bellani M, Liu Y, Seidman MM. Fanconi Anemia: a DNA Repair Disorder Characterized by Accelerated Decline of the Hematopoietic Stem Cell Compartment and Other Features of Aging. Ageing Res Rev [Internet]. NIH Public Access; 2017 [cited 2024 Oct 15];33:67. 10.1016/J.ARR.2016.05.005

14. Lum SH, Eikema DJ, Piepenbroek B, Wynn RF, Samarasinghe S, Dalissier A, et al. Outcomes of hematopoietic stem cell transplantation in 813 pediatric patients with Fanconi anemia. Blood. Elsevier B.V.; 2024;144:1329–42. 10.1182/BLOOD.2023022751

15. Dufour C, Pierri F. Modern management of Fanconi anemia. Hematology Am Soc Hematol Educ Program [Internet]. Hematology Am Soc Hematol Educ Program; 2022;2022:649–57. 10.1182/HEMATOLOGY.2022000393

16. Cancio M, Lucas AGT, Bierings M, Klein E, de Witte MA, Smiers FJ, et al. Predictors of outcomes in hematopoietic cell transplantation for Fanconi anemia. Bone Marrow Transplant. Springer Nature; 2024;59:34–40. 10.1038/S41409-023-02121-1

17. Hoover A, Turcotte LM, Phelan R, Barbus C, Rayannavar A, Miller BS, et al. Longitudinal clinical manifestations of Fanconi anemia: A systematized review. Blood Rev. Churchill Livingstone; 2024;68. 10.1016/J.BLRE.2024.101225

18. Fink O, Even-Or E, Avni B, Grisariu S, Zaidman I, Schejter YD, et al. Two decades of stem cell transplantation in patients with Fanconi anemia: Analysis of factors affecting transplant outcomes. Clin Transplant. John Wiley and Sons Inc; 2023;37. 10.1111/CTR.14835

19. Ricci E, Bagnasco F, Pierri F, Risitano A, Farruggia P, Faraci M, et al. Long-Term Outcome of Fanconi Anemia Patients From the Italian Registry on Behalf of the Marrow Failure Study Group of the AIEOP (Italian Association for Pediatric Haematology-Oncology). Am J Hematol. John Wiley and Sons Inc; 2025;100:1387–96. 10.1002/AJH.27724

20. Morrison SJ, Scadden DT. The bone marrow niche for haematopoietic stem cells. Nature [Internet]. Nature; 2014;505:327–34. 10.1038/NATURE12984

21. Mendelson A, Frenette PS. Hematopoietic stem cell niche maintenance during homeostasis and regeneration. Nat Med [Internet]. Nat Med; 2014;20:833–46. 10.1038/NM.3647

22. Park M. Overview of inherited bone marrow failure syndromes. Blood Res. Korean Society of Hematology; 2022;57:49–54. 10.5045/BR.2022.2022012

23. Kallen ME, Dulau-Florea A, Wang W, Calvo KR. Acquired and germline predisposition to bone marrow failure: Diagnostic features and clinical implications. Semin Hematol. W.B. Saunders; 2019;56:69–82. 10.1053/J.SEMINHEMATOL.2018.05.016

24. Frenette PS, Méndez-Ferrer S, Lucas-Alcaraz D, Batista M, Lira S, Michurina T V, et al. The Hematopoietic Stem Cell Niche. Blood [Internet]. 2009;114:SCI-49-SCI-49. 10.1182/BLOOD.V114.22.SCI-49.SCI-49

25. Celso C Lo, Scadden DT. The haematopoietic stem cell niche at a glance. J Cell Sci. 2011;124:3529–35. 10.1242/JCS.074112

26. Frenette P. Hematopoietic stem cell niche through the ages. Exp Hematol. Elsevier BV; 2016;44:S33. 10.1016/J.EXPHEM.2016.06.027

27. Testa U, Castelli G, Pelosi E. Curative Approach to the Treatment of Beta-Thalassemia and Sickle Cell Disease with Hematopoietic Stem Cell Transplantation. J Clin Med. Multidisciplinary Digital Publishing Institute (MDPI); 2026;15. 10.3390/JCM15041379

28. Neoman N, Kim HN, Viduya J, Goyal A, Liu YL, Sakamoto KM. Inflammatory pathways and the bone marrow microenvironment in inherited bone marrow failure syndromes. Stem Cells. Oxford University Press; 2025;43. 10.1093/STMCLS/SXAF021

29. Kawashima N, Bezzerri V, Corey SJ. The Molecular and Genetic Mechanisms of Inherited Bone Marrow Failure Syndromes: The Role of Inflammatory Cytokines in Their Pathogenesis. Biomolecules. Multidisciplinary Digital Publishing Institute (MDPI); 2023;13. 10.3390/BIOM13081249

30. Balbi M, Guidi E, Hristodor AM, Corsolini F, Cossu V, Bottega R, et al. Beyond readthrough: ataluren restores mitochondrial function and reduces oxidative stress in FANCA-mutated cells via mTOR–DRP1 modulation. Cell Death Discovery 2026 [Internet]. Nature Publishing Group; 2026; 10.1038/s41420-026-02983-6

31. Raggi F, Bartolucci M, Cangelosi D, Rossi C, Pelassa S, Trincianti C, et al. Proteomic profiling of extracellular vesicles in synovial fluid and plasma from Oligoarticular Juvenile Idiopathic Arthritis patients reveals novel immunopathogenic biomarkers. Front Immunol [Internet]. Front Immunol; 2023;14. 10.3389/FIMMU.2023.1134747

32. Ravera S, Podestà M, Sabatini F, Dagnino M, Cilloni D, Fiorini S, et al. Discrete Changes in Glucose Metabolism Define Aging. Sci Rep [Internet]. Nature Publishing Group; 2019 [cited 2022 Mar 15];9:10347. 10.1038/s41598-019-46749-w

33. Cossu V, Bertola N, Fresia C, Sabatini F, Ravera S. Redox Imbalance and Antioxidant Defenses Dysfunction: Key Contributors to Early Aging in Childhood Cancer Survivors. Antioxidants (Basel) [Internet]. Antioxidants (Basel); 2024;13. 10.3390/ANTIOX13111397

34. Ding W, Kaur D, Horvath S, Zhou W. Comparative epigenome analysis using Infinium DNA methylation BeadChips. Brief Bioinform [Internet]. Brief Bioinform; 2023;24. 10.1093/BIB/BBAC617

35. Du P, Zhang X, Huang CC, Jafari N, Kibbe WA, Hou L, et al. Comparison of Beta-value and M-value methods for quantifying methylation levels by microarray analysis. BMC Bioinformatics [Internet]. BioMed Central; 2010;11:587. 10.1186/1471-2105-11-587/FIGURES/5

36. Johnson WE, Li C, Rabinovic A. Adjusting batch effects in microarray expression data using empirical Bayes methods. Biostatistics [Internet]. Biostatistics; 2007;8:118–27. 10.1093/BIOSTATISTICS/KXJ037

37. Marioni RE, Hillary RF. MethylDetectR: A software for methylation-based health profiling. Wellcome Open Res [Internet]. F1000 Research Ltd; 2021;5. 10.12688/WELLCOMEOPENRES.16458.2/DOI

38. Cappelli E, Ravera S, Vaccaro D, Cuccarolo P, Bartolucci M, Panfoli I, et al. Mitochondrial respiratory complex I defects in Fanconi anemia. Trends Mol Med. 2013;19. 10.1016/j.molmed.2013.07.008

39. Cappelli E, Bertola N, Bruno S, Degan P, Regis S, Corsolini F, et al. A Multidrug Approach to Modulate the Mitochondrial Metabolism Impairment and Relative Oxidative Stress in Fanconi Anemia Complementation Group A. Metabolites [Internet]. Multidisciplinary Digital Publishing Institute; 2021;12:6. 10.3390/metabo12010006

40. Bertola N, Regis S, Bruno S, Mazzarello AN, Serra M, Lupia M, et al. Effects of Deacetylase Inhibition on the Activation of the Antioxidant Response and Aerobic Metabolism in Cellular Models of Fanconi Anemia. Antioxidants (Basel) [Internet]. Antioxidants (Basel); 2023 [cited 2024 Feb 13];12. 10.3390/ANTIOX12051100

41. Bellanti F, Coda ARD, Trecca MI, Buglio A Lo, Serviddio G, Vendemiale G. Redox Imbalance in Inflammation: The Interplay of Oxidative and Reductive Stress. Antioxidants [Internet]. Multidisciplinary Digital Publishing Institute (MDPI); 2025;14:656. 10.3390/ANTIOX14060656

42. Lei Y, Wang K, Deng L, Chen Y, Nice EC, Huang C. Redox Regulation of Inflammation: Old Elements, a New Story. Med Res Rev [Internet]. John Wiley & Sons, Ltd; 2015;35:306–40. 10.1002/MED.21330

43. Pagliara D, Ciolfi A, Pedace L, Haghshenas S, Ferilli M, Levy MA, et al. Identification of a robust DNA methylation signature for Fanconi anemia. Am J Hum Genet [Internet]. Am J Hum Genet; 2023;110:1938– 49. 10.1016/J.AJHG.2023.09.014

44. Dufour C, Corcione A, Svahn J, Haupt R, Poggi V, Béka’ssy AN, et al. TNF-alpha and IFN-gamma are overexpressed in the bone marrow of Fanconi anemia patients and TNF-alpha suppresses erythropoiesis in vitro. Blood [Internet]. 2003;102:2053–9. 10.1182/blood-2003-01-0114

45. Mathavarajah S, Dellaire G. LINE-1: an emerging initiator of cGAS-STING signalling and inflammation that is dysregulated in disease. Biochemistry and Cell Biology. Canadian Science Publishing; 2024;102:38–46. 10.1139/BCB-2023-0134

46. Bertola N, Bruno S, Capanni C, Columbaro M, Mazzarello AN, Corsolini F, et al. Altered Mitochondrial Dynamic in Lymphoblasts and Fibroblasts Mutated for FANCA-A Gene: The Central Role of DRP1. International Journal of Molecular Sciences 2023, Vol 24, Page 6557 [Internet]. Multidisciplinary Digital Publishing Institute; 2023 [cited 2023 Apr 27];24:6557. 10.3390/IJMS24076557

47. Velleuer E, Carlberg C. Fanconi anaemia as a human model of accelerated epigenetic and immune ageing. Ageing Res Rev. Elsevier; 2026;115:103038. 10.1016/J.ARR.2026.103038

48. Degan P, Cappelli E, Regis S, Ravera S. New Insights and Perspectives in Fanconi Anemia Research. Trends Mol Med. 2019;25. 10.1016/j.molmed.2019.01.003

49. Ayala A, Muñoz MF, Argüelles S. Lipid peroxidation: production, metabolism, and signaling mechanisms of malondialdehyde and 4-hydroxy-2-nonenal. Oxid Med Cell Longev [Internet]. 2014;2014:360438. 10.1155/2014/360438

50. Uchida K. 4-Hydroxy-2-nonenal: A product and mediator of oxidative stress. Prog Lipid Res. Elsevier Ltd; 2003;42:318–43. 10.1016/S0163-7827(03)00014-6

51. Poli G, Schaur RJ, Siems WG, Leonarduzzi G. 4-Hydroxynonenal: A membrane lipid oxidation product of medicinal interest. Med Res Rev. 2008;28:569–631. 10.1002/MED.20117

52. Radi R. Protein tyrosine nitration: Biochemical mechanisms and structural basis of functional effects. Acc Chem Res. 2013;46:550–9. 10.1021/AR300234C

53. Du W, Rani R, Sipple J, Schick J, Myers KC, Mehta P, et al. The FA pathway counteracts oxidative stress through selective protection of antioxidant defense gene promoters. Blood [Internet]. American Society of Hematology; 2012;119:4142. 10.1182/BLOOD-2011-09-381970

54. Zhou Y, He Y, Xing W, Zhang P, Shi H, Chen S, et al. An abnormal bone marrow microenvironment contributes to hematopoietic dysfunction in fanconi anemia. Haematologica. Ferrata Storti Foundation; 2017;102:1017–27. 10.3324/HAEMATOL.2016.158717

55. Du W, Erden O, Pang Q. TNF-α signaling in Fanconi anemia. Blood Cells Mol Dis [Internet]. Blood Cells Mol Dis; 2014;52:2–11. 10.1016/J.BCMD.2013.06.005

56. Korthof ET, Svahn J, Peffault de Latour R, Terranova P, Moins-Teisserenc H, Socié G, et al. Immunological profile of Fanconi anemia: a multicentric retrospective analysis of 61 patients. Am J Hematol [Internet]. 2013;88:472–6. 10.1002/ajh.23435

57. Río P, Bueren JA. TGF-β: A master regulator of the bone marrow failure puzzle in Fanconi anemia. Stem Cell Investig. AME Publishing Company; 2016;2016. 10.21037/SCI.2016.09.17

58. Pastrez PRA, Barbosa AM, Mariano VS, Causin RL, Castro AG, Torrado E, et al. Interleukin-8 and Interleukin-6 Are Biomarkers of Poor Prognosis in Esophageal Squamous Cell Carcinoma. Cancers (Basel) [Internet]. MDPI; 2023;15:1997. 10.3390/CANCERS15071997/S1

59. Anur P, Friedman DN, Sklar C, Oeffinger K, Castiel M, Kearney J, et al. Late effects in patients with Fanconi anemia following allogeneic hematopoietic stem cell transplantation from alternative donors. Bone Marrow Transplant. Nature Publishing Group; 2016;51:938–44. 10.1038/BMT.2016.32

60. Zhang X, Chen X, Song H, Chen HZ, Rovin BH. Activation of the Nrf2/antioxidant response pathway increases IL-8 expression. Eur J Immunol [Internet]. Eur J Immunol; 2005;35:3258–67. 10.1002/EJI.200526116

61. Trino S, Zoppoli P, Carella AM, Laurenzana I, Weisz A, Memoli D, et al. DNA methylation dynamic of bone marrow hematopoietic stem cells after allogeneic transplantation. Stem Cell Res Ther [Internet]. BioMed Central Ltd.; 2019;10:138. 10.1186/S13287-019-1245-6/FIGURES/6

62. Hristodor AM, Cappelli E, Baldisseri E, Valli R, Montalbano G, Micheloni G, et al. Development of translational read-through-inducing drugs as novel therapeutic options for patients with Fanconi anemia. Cell Death Discovery 2025 11:1 [Internet]. Nature Publishing Group; 2025;11:1–13. 10.1038/s41420-025-02571-0

