## Supplementary Figures for "Recipient-driven metabolic and epigenetic remodeling of donor hematopoietic cells after transplantation in Fanconi Anemia"

#### The Bone Marrow Microenvironment in Transplanted Fanconi Anemia Patients and its Influence on Donor Cells

[d] Unit of Hematology, IRCCS Istituto Giannina Gaslini, Genoa, Italy.

[e] Hematopoietic Stem Cell Transplantation Unit, Department of Pediatric Hematology-Oncology, IRCCS Istituto Giannina Gaslini, Genoa, Italy.

[f] Diagnosis, Observation, and Prevention after Oncological therapy (D.O.P.O.) Clinic, Department of Pediatric Hematology and Oncology, Istituto di Ricovero e Cura a Carattere Scientifico (IRCCS), Istituto Giannina Gaslini, Genoa, Italy.

[g] Clinical and Experimental Immunology Unit, Department of Services, IRCCS Istituto Giannina Gaslini, Genoa, Italy.

[h] AOM - IRCCS Ospedale Policlinico San Martino, Genoa, Italy

[§] These authors contributed equally as first authors

[#] These authors contributed equally as senior authors.

* Corresponding Author:

Prof. Silvia Ravera

University of Genoa, Department of Experimental Medicine

Via De Toni 14, 16132 Genoa, Italy

**Supplementary Materials and Methods**

**1. EVs isolation from HD and FA PB**

Extracellular vesicles (EVs) were isolated from 500 µl of plasma (PL) samples using the exoRNeasy Serum/Plasma Midi Kit (Qiagen Italia, Milan, Italy), which employs membrane affinity spin columns to capture intact EVs from small sample volumes efficiently, according to the manufacturer's instructions and as previously described [1]. Briefly, plasma samples were centrifuged at 500 × g for 10 min at 25 °C, followed by centrifugation at 16,000 × g for 15 min to remove cells and cellular debris. The resulting supernatants were mixed with an equal volume of XBP Binding Buffer and loaded onto exoEasy spin columns to allow EV binding to the membrane. After centrifugation at 500 × g for 1 min at RT, the columns were washed with 3.5 mm of XWP Washing Buffer and centrifuged at 5,000 × g for 5 min at RT to remove non-specifically bound material. The spin columns were then transferred to clean collection tubes, and 100 µl of XE Buffer was added directly onto the membrane. After a 1-min incubation at RT, EVs were eluted by centrifugation at 500 × g for 5 min. The eluate was subsequently reloaded onto the same spin column and centrifuged again at 5,000 × g for 5 min to maximize EV recovery.

**2. Cell homogenate preparation and protein quantification**

Cells were collected by centrifugation, washed with PBS, and resuspended in PBS supplemented with protease and phosphatase inhibitors. Cell suspensions were sonicated on ice for two cycles of 10 s with a 30 s interval. Protein concentration was determined by Bradford assay, using bovine serum albumin as standard [2].

**3. ATP synthesis, oxygen consumption rate, and P/O ratio**

OxPhos was evaluated by measuring ATP synthesis, oxygen consumption rate (OCR), and coupling efficiency expressed as the P/O ratio.

For each experiment, 50,000 cells were permeabilized with digitonin 0.03 mg/ml for 1 min before analysis, or $10^8$ EVs were employed.

ATP synthesis was quantified at 25°C using a luciferin/luciferase ATP bioluminescence assay (Roche Diagnostics Corp., Indianapolis, IN, USA). Samples were resuspended in a reaction medium containing PBS plus 0.6 mM ouabain, 0.2 mM P1,P5-Di(adenosine-5') pentaphosphate, and 10 mM pyruvate plus 5 mM malate (P/M) or20 mM succinate (Succ), as respiring substrates to evaluate complex I-linked or complex II-linked respiration, respectively. ATP synthesis was initiated by addition of 0.1 mM ADP and monitored every 30 seconds for 2 min by a luminometer (GloMax® 20/20n Luminometer, Promega Italia, Milano, Italy). ATP concentrations were calculated from a standard curve generated with ATP standards with a range between $10^{-10}$–$10^{-7}$ M [3].

OCR was measured at 25°C using an amperometric oxygen electrode (Unisense Microrespiration, Unisense A/S, Aarhus, Denmark) in a sealed respiration chamber. Samples were suspended in PBS and permeabilized with digitonin 0.03 mg/ml for 1 min before analysis. Respiration was stimulated with the same respiratory substrates used for ATP synthesis. ADP-dependent respiration was induced by adding 0.1mM ADP [3].

OxPhos efficiency was calculated as the P/O ratio, defined as the amount of ATP produced relative to the amount of oxygen consumed over the same time interval. Values close to 2.5 for P/M and 1.5 for Succ were considered compatible with efficient coupling, whereas lower values were interpreted as reduced coupling efficiency [4].

**4. Intracellular ATP, AMP, and ATP/AMP ratio**

Intracellular ATP and AMP levels were quantified spectrophotometrically on 50 µg of total protein, following enzyme-coupled reactions [3].

ATP was measured by monitoring the reduction of NADP at 340 nm in a reaction mixture containing 100 mM Tris-HCl (pH 8.0), 0.2 mM NADP, 5 mM MgCl2, and 50 mM glucose. Samples were analysed before and after the addition of 3 µg of purified hexokinase plus glucose-6-phosphate dehydrogenase.

AMP was measured by monitoring NADH oxidation at 340 nm in the presence of 100 mM Tris-HCl (pH 8.0), 5 mM MgCl2, 0.2 mM ATP, 10 mM phosphoenolpyruvate, 0.15 mM NADH, 10 IU adenylate kinase, 25 IU pyruvate kinase, and 15 IU lactate dehydrogenase.

Cellular energy status was expressed as the ATP/AMP ratio.

**5. Assessment of oxidative damage markers**

Lipid peroxidation was evaluated by measuring malondialdehyde (MDA) using the thiobarbituric acid reactive substances (TBARS) assay. Briefly, 50 µg of total protein of cell homogenate or $10^8$ EVs were diluted in 300 µL of Milli-Q water and mixed with 600 µL of TBARS reagent containing trichloroacetic acid, hydrochloric acid, and thiobarbituric acid. Samples were incubated at 95 °C for 60 min, centrifuged, and absorbance was read at 532 nm. MDA concentration was calculated using a calibration curve generated with MDA standards ranging from 1 to 20 µM [3]. Comparable TBARS-based assays were used in the attached FA and redox-metabolism studies.

4-hydroxynonenal (4-HNE, #ab287803), nitrotyrosine (#ab116691), and 8-hydroxy-2′-deoxyguanosine (8-OHdG, #ab201734) were measured using commercially available ELISA kits (Abcam, Cambridge, UK).

**6. Glutathione reductase and catalase activity**

Glutathione reductase (GR) and catalase (CAT) activities were measured spectrophotometrically in cell homogenates. For each assay, 50 µg of total protein were used.

GR activity was assessed by monitoring NADPH oxidation at 340 nm in a reaction mixture containing 100 mM Tris-HCl (pH 7.4), 1 mM EDTA, 5 mM GSSG, and 0.2 mM NADPH [5].

CAT activity was measured by following hydrogen peroxide decomposition at 240 nm in a reaction mixture containing 50 mM phosphate buffer (pH 7.0) and 5 mM $H_2O_2$ [5].

**7. Cytokine quantification**

TGF-β1 was quantified in peripheral blood (PB) plasma using the Simple Plex Human TGF-beta 1 Cartridge (# SPCKB-PS-001369), whereas TNF-α, IL-6, and IL-8 were quantified using the Simple Plex Cytokine Storm Panel (# ST01A-PS-003229) on the Ella automated immunoassay platform, according to the manufacturer's instructions (Bio-Techne, Minneapolis, MN, USA). Cytokine concentrations were calculated from standard curves and expressed as pg/ml.

**8. Western blot analysis and densitometry**

Western blot (WB) analysis was performed on cell lysates prepared from PB-MNCs. Equal amounts of protein, typically 30 µg per lane, were separated by SDS-PAGE using 4–20% gradient gels (Bio-Rad, Hercules, CA, USA) and transferred onto nitrocellulose membranes.

Membranes were blocked and incubated overnight at 4 °C with primary antibodies against NRF2 (#12721), phosphorylated STAT2 (pSTAT2, #88410), total STAT2 (#72604), phosphorylated SMAD3 (pSMAD3, #9520), total SMAD3 (#9523), and actin (#3700). All antibodies were from Cell Signalling Technologies and were diluted 1:200 in PBS plus 0.15% Tween (PBSt).

After washing with PBSt, Secondary antibodies (#A0168 and #SAB3700870, Sigma-Aldrich, St. Louis, MO, USA) were diluted 1:10,000 in PBSt. Bands were evaluated by a chemiluminescence system (Alliance 6.7 WL 20M, UVITEC, Cambridge, UK) using an enhanced chemiluminescence substrate (ECL, #1705061, BioRad, Hercules, CA, USA). Band intensity was evaluated by Alliance TM Q9-Series software (UVITEC, Cambridge, UK). All the signals were normalized versus the actin signal.

**9. DNA methylation and epigenetic analyses**

Genomic DNA was extracted from peripheral blood mononuclear cells (PBMC) using High Pure PCR Template Preparation Kit (Roche Diagnostics, Mannheim, Germany). DNA concentrations were assessed by Qubit™ dsDNA Quantification Assay Kit (Thermo Fisher Scientific, Waltham, MA, USA). Genome-wide DNAm profiling was conducted using the Illumina Methylation Screening Array (MSA; Illumina, San Diego, CA, USA) at a certified external service provider (Helmholtz Munich, Germany), following the manufacturer's protocol.

Raw intensity data were pre-processed following a well-established pipeline implemented in the *sesame* R package using the *openSesame* function, which includes background correction, dye-bias adjustment, probe-type normalization, removal of cross-reactive probes and those mapping within a single nucleotide polymorphism (SNPs), following quality control filtering of low-quality probes and samples [6]. DNAm levels were quantified as β values, defined as the ratio of methylated probe intensity to the sum of methylated and unmethylated probe intensities. For downstream statistical analyses, a logit transformation of β values was applied to derive M values, defined as the log2 ratio of methylated to unmethylated probe intensities [7]. Batch effect correction was performed to remove unwanted technical variation arising from differences in processing batches using the ComBat function within the sva R package [8]. Principal Component Analysis (PCA) was performed in R statistical software v.4.2.2. Additionally, we computed epigenetic scores for 109 circulating proteins using the *MethylDetectR* R package [9]. Briefly, MethylDetectR estimates protein levels from DNA methylation data by applying previously validated CpG-based predictive models trained on large population cohorts, thereby providing methylation-derived proxies (episcores) for circulating protein abundance. These episcores were used to investigate whether the persistent inflammatory phenotype observed after HSCT may be attributable, at least in part, to an epigenetic memory that influences the regulation of proteins involved in biological pathways relevant to the FA recipient microenvironment.

**References**

1. Raggi F, Bartolucci M, Cangelosi D, et al. Proteomic profiling of extracellular vesicles in synovial fluid and plasma from Oligoarticular Juvenile Idiopathic Arthritis patients reveals novel immunopathogenic biomarkers. *Front Immunol*. 2023;14. doi:10.3389/FIMMU.2023.1134747
2. Bradford MM. A rapid and sensitive method for the quantitation of microgram quantities of protein utilizing the principle of protein-dye binding. *Anal Biochem*. 1976;72(1-2):248-254. doi:10.1016/0003-2697(76)90527-3
3. Cappelli E, Degan P, Bruno S, et al. The passage from bone marrow niche to bloodstream triggers the metabolic impairment in Fanconi Anemia mononuclear cells. *Redox Biol*. 2020;36. doi:10.1016/j.redox.2020.101618
4. Ravera S, Podestà M, Sabatini F, et al. Discrete Changes in Glucose Metabolism Define Aging. *Sci Rep*. 2019;9(1):10347. doi:10.1038/s41598-019-46749-w
5. Cossu V, Bertola N, Fresia C, Sabatini F, Ravera S. Redox Imbalance and Antioxidant Defenses Dysfunction: Key Contributors to Early Aging in Childhood Cancer Survivors. *Antioxidants (Basel)*. 2024;13(11). doi:10.3390/ANTIOX13111397

6. Ding W, Kaur D, Horvath S, Zhou W. Comparative epigenome analysis using Infinium DNA methylation BeadChips. *Brief Bioinform*. 2023;24(1). doi:10.1093/BIB/BBAC617
7. Du P, Zhang X, Huang CC, et al. Comparison of Beta-value and M-value methods for quantifying methylation levels by microarray analysis. *BMC Bioinformatics*. 2010;11(1):587. doi:10.1186/1471-2105-11-587/FIGURES/5
8. Johnson WE, Li C, Rabinovic A. Adjusting batch effects in microarray expression data using empirical Bayes methods. *Biostatistics*. 2007;8(1):118-127. doi:10.1093/BIOSTATISTICS/KXJ037
9. Marioni RE, Hillary RF. MethylDetectR: A software for methylation-based health profiling. *Wellcome Open Res*. 2021;5. doi:10.12688/WELLCOMEOPENRES.16458.2/DOI

### Supplementary Figure 1

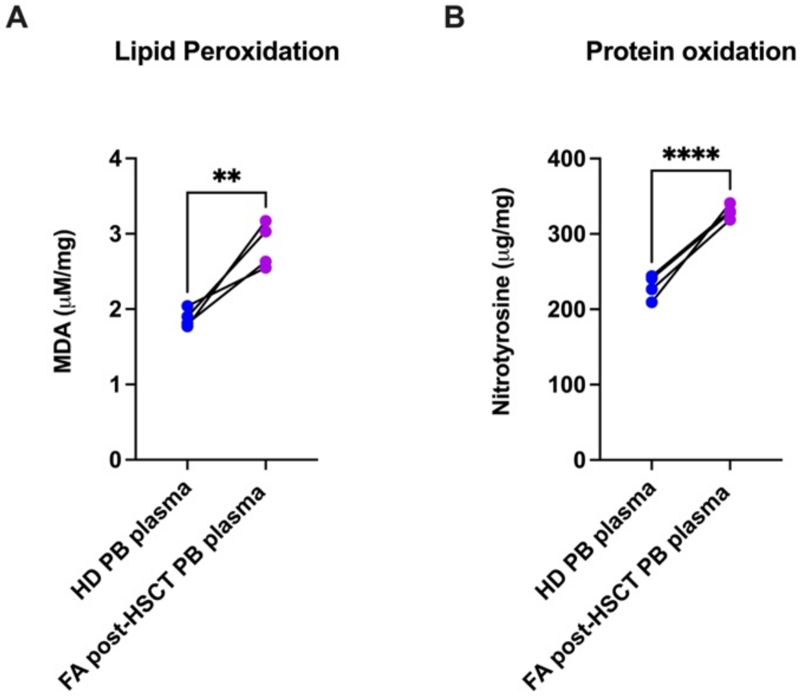


**Supplementary Figure 1. Paired analysis of oxidative stress markers in donors and FA recipients after HCT.**
MDA (A) and nitrotyrosine (B) levels were measured in peripheral blood (PB) plasma from healthy donors (HD) and the corresponding patients with Fanconi anaemia after HCT (FA post-HCT). Each dot represents an individual sample, and lines connect each donor with the corresponding recipient (n = 4 donor–recipient pairs). Statistical analysis was performed using a paired Student’s t-test. ****$p < 0.0001$.

### Supplementary Figure 2

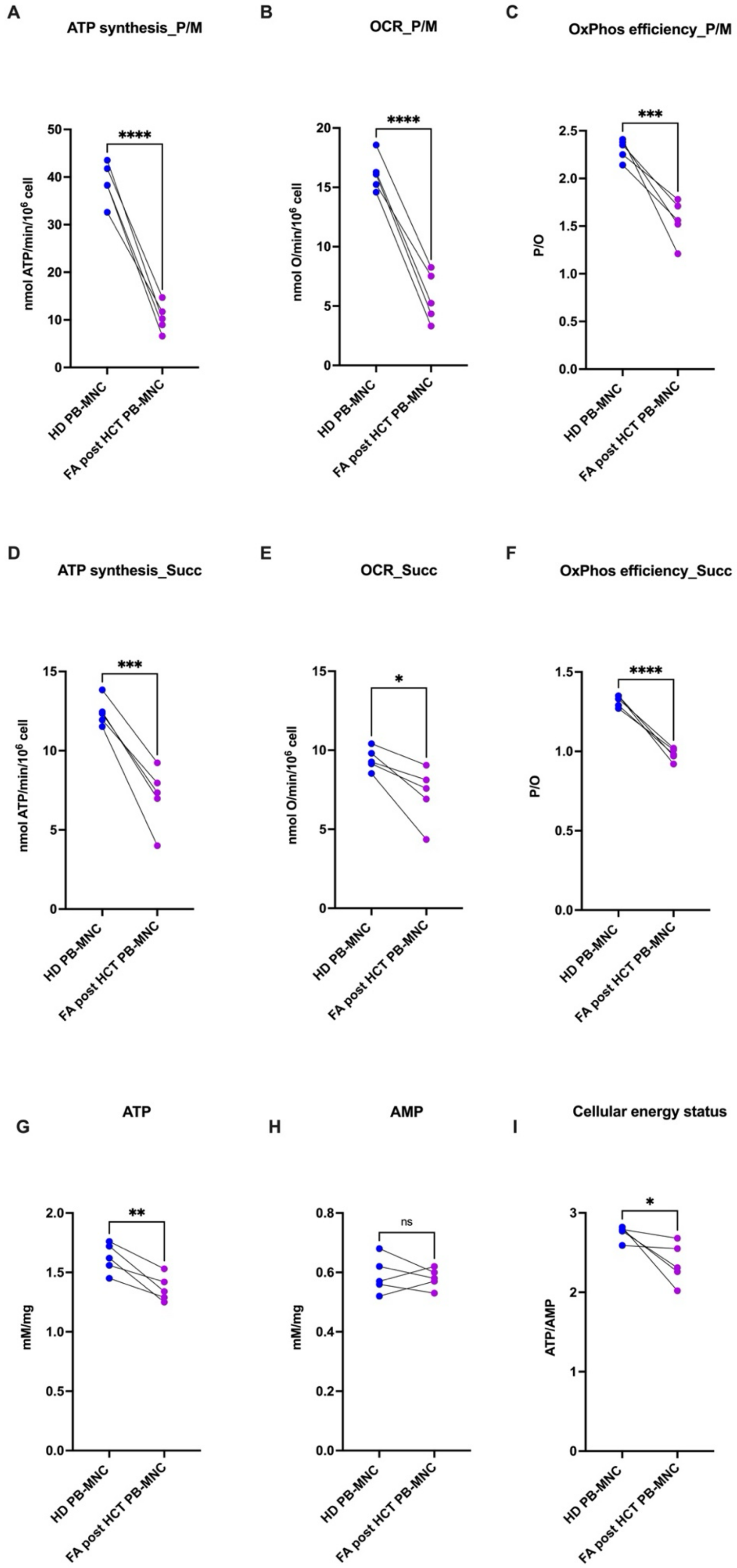

**Supplementary Figure 2. Paired analysis of mitochondrial respiration and cellular energy status in donors and FA recipients after HCT.** Mitochondrial respiration and cellular energy status were measured in peripheral blood (PB) plasma from healthy donors (HD) and the corresponding patients with Fanconi anaemia after HCT (FA post-HCT). (A–C) ATP synthesis, oxygen consumption rate (OCR), and OxPhos efficiency, expressed as P/O ratio, were measured under pyruvate/malate-driven (P/M) respiration. (D–F) ATP synthesis, OCR, and OxPhos efficiency, expressed as P/O ratio, were measured under succinate-driven (Succ) respiration. (G) Intracellular ATP levels. (H)Intracellular AMP levels. (I)Cellular energy status expressed as the ATP/AMP ratio. Each dot represents an individual sample, and lines connect each donor with the corresponding recipient (n = 5 donor–recipient pairs). Statistical analysis was performed using a paired Student's t-test. ns, not significant; *$p < 0.05$, **$p < 0.01$, ***$p < 0.001$, ****$p < 0.0001$

**Supplementary Figure 3**

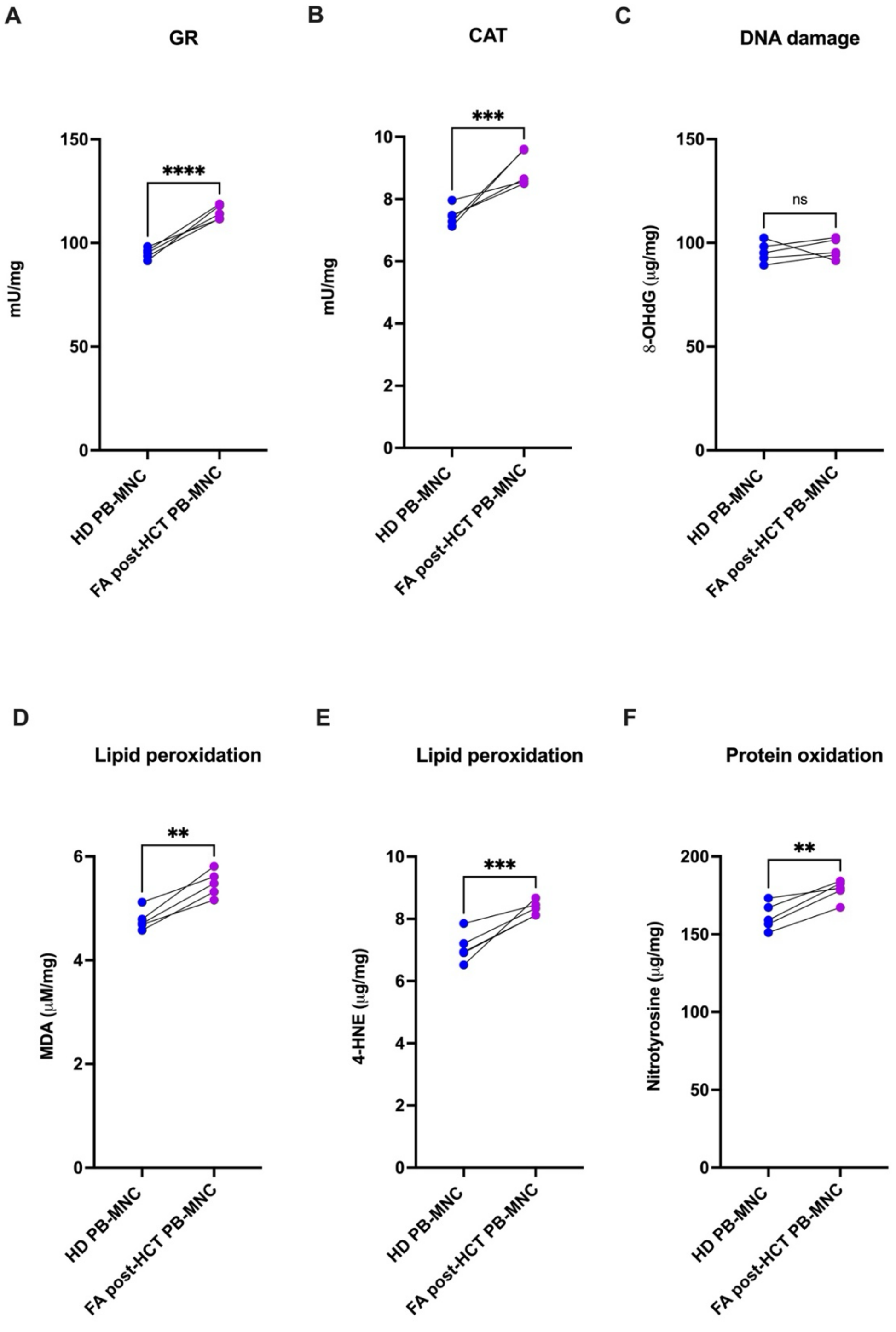


**Supplementary Figure 3. Paired analysis of oxidative damage markers and antioxidant enzyme activities in donors and FA recipients after HCT.**

Mitochondrial respiration and cellular energy status were measured in peripheral blood (PB) mononuclear cells (MNCs) from healthy donors (HD) and the corresponding patients with Fanconi anaemia after HCT (FA post-HCT).(A) Glutathione reductase (GR) activity. (B)Catalase (CAT) activity. (C) DNA oxidative damage assessed as 8-hydroxy-2′-deoxyguanosine (8-OHdG) levels. (D, E) Lipid peroxidation, assessed as malondialdehyde (MDA) and 4-

hydroxynonenal (4-HNE) levels, respectively. (F) Protein oxidation, assessed as nitrotyrosine levels. Each dot represents an individual sample, and lines connect each donor with the corresponding recipient (n = 5 donor–recipient pairs). Statistical analysis was performed using a paired Student’s t-test. ns, not significant; $^{**}p < 0.01$, $^{***}p < 0.001$, $^{****}p < 0.0001$.

### Supplementary Figure 4

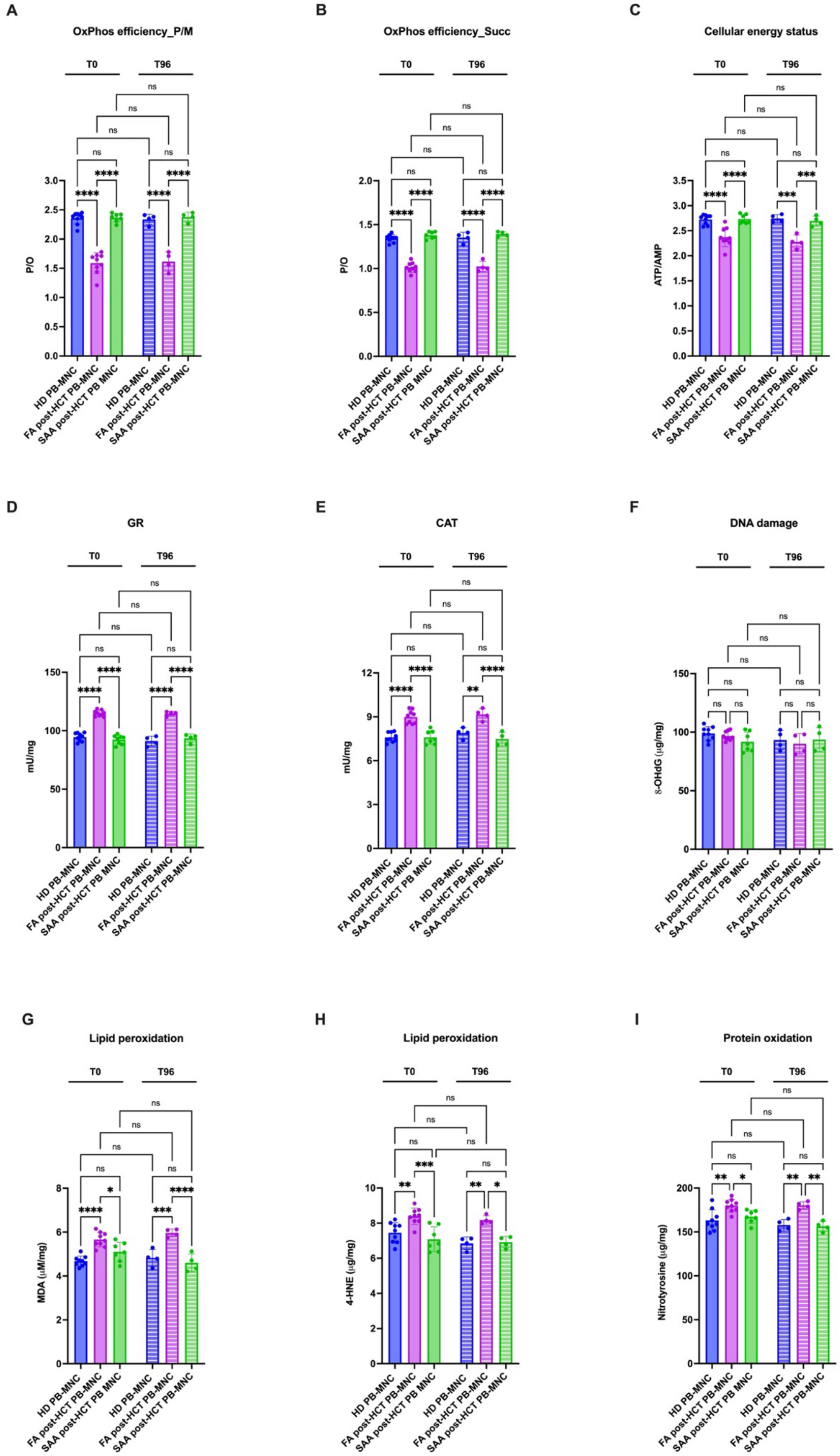

**Supplementary Figure 4. Persistent metabolic and redox alterations in PB-MNCs from FA patients after HCT following 96 h of culture.**

Peripheral blood mononuclear cells from healthy donors (HD PB-MNC), non-transplanted Fanconi Anemia (FA PB-MNC) patients, FA patients after hematopoietic stem cell transplantation (FA post-HCT PB-MNC), and Severe Aplastic Anemia patients after hematopoietic stem cell transplantation (SAA post-HCT PB-MNC) were analyzed for antioxidant enzyme activity and oxidative damage markers. (A)OxPhos efficiency, expressed as P/O ratio measured under pyruvate/malate-driven (P/M) respiration. (B) OxPhos efficiency, expressed as P/O ratio measured under succinate-driven (Succ) respiration. (C) Cellular energy status expressed as the ATP/AMP ratio. (D) Glutathione reductase (GR) activity. (E) Catalase (CAT) activity. (F) DNA oxidative damage, assessed as 8-hydroxy-2′-deoxyguanosine (8-OHdG) levels. (G, H) Lipid peroxidation, assessed as malondialdehyde (MDA) and 4-hydroxynonenal (4-HNE) levels, respectively. (I) Protein oxidation, assessed as nitrotyrosine levels. Data are presented as mean ± SD ($n \geq 4$ for each panel). Statistical analysis was performed using one-way analysis of variance (ANOVA), followed by Tukey's post hoc multiple comparison test. ns, not significant; *$p < 0.05$, **$p < 0.01$, ***$p < 0.001$, ****$p < 0.0001$.

### Supplementary Figure 5

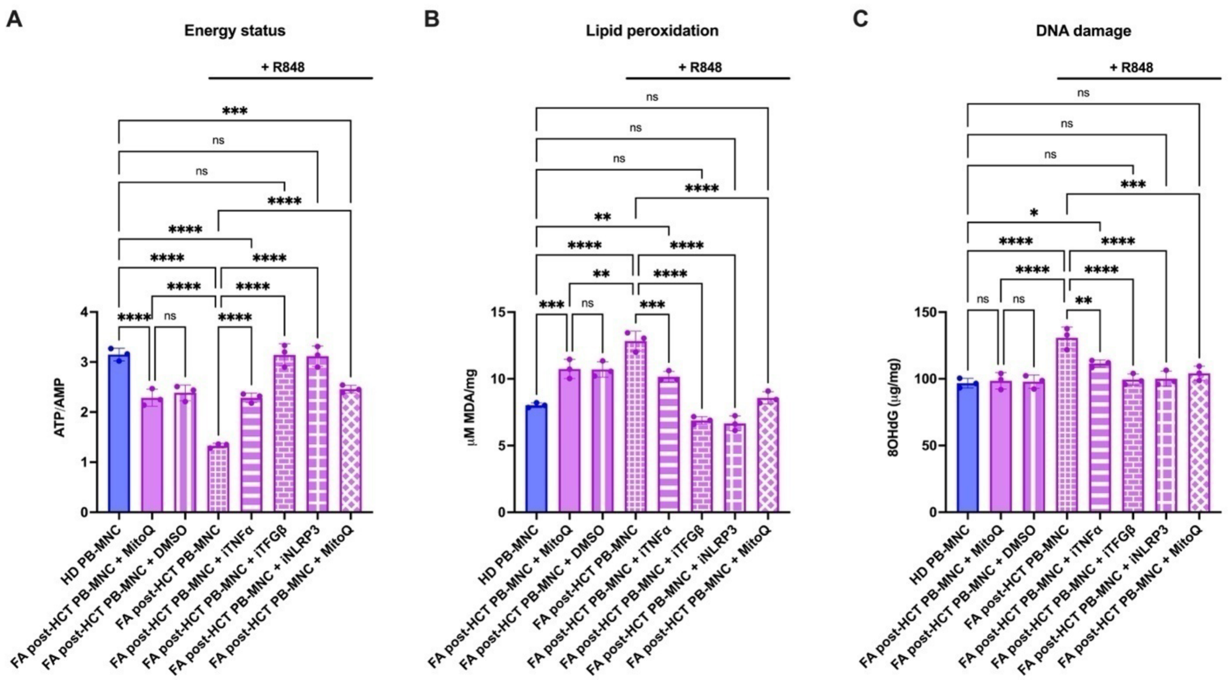


**Supplementary Figure 5. Effect of inflammatory-pathway inhibition and mitochondrial antioxidant treatment on R848-stimulated FA post-HCT PB-MNCs.**

HD PB-MNCs and FA post-HCT PB-MNCs were analysed under basal conditions. FA post-HCT PB-MNCs were also stimulated with DMSO, as a vehicle control, and R848, alone or in combination with inhibitors targeting TNF-α, TGF-β, and NLRP3-related pathways, or with the mitochondria-targeted antioxidant MitoQ.(A) Cellular energy status expressed as ATP/AMP ratio.(B) Lipid peroxidation assessed by MDA levels.(C) Oxidative DNA damage assessed by 8OHdG levels. Data are presented as mean ± SD (n = 3 for each panel). Statistical analysis was performed using one-way analysis of variance (ANOVA), followed by Tukey's post hoc multiple comparison test. ns, not significant; **p < 0.01, *** p < 0.001, ****p < 0.0001.
