## Supplementary figures and images for "Recipient-driven metabolic and epigenetic remodeling of donor hematopoietic cells after transplantation in Fanconi Anemia"

### Whole WB signals

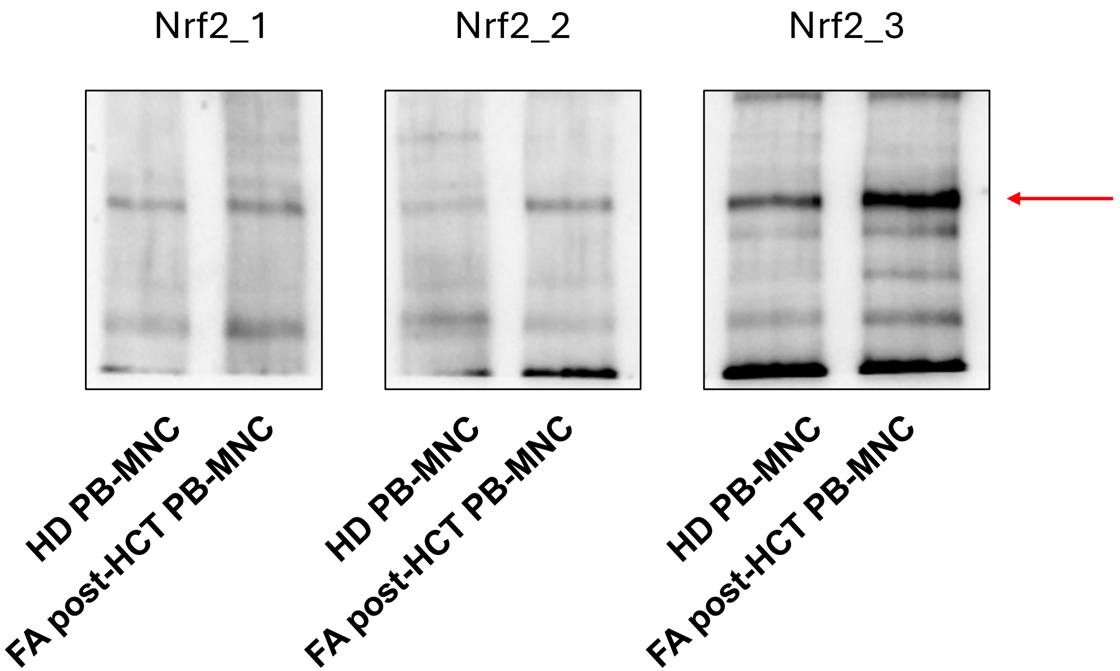

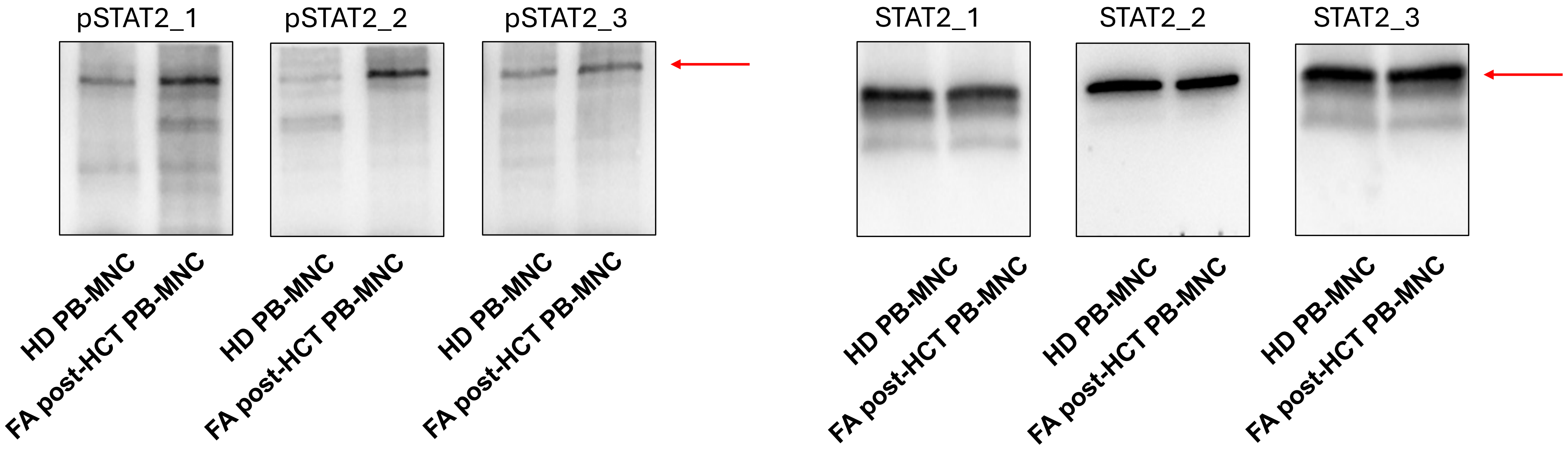

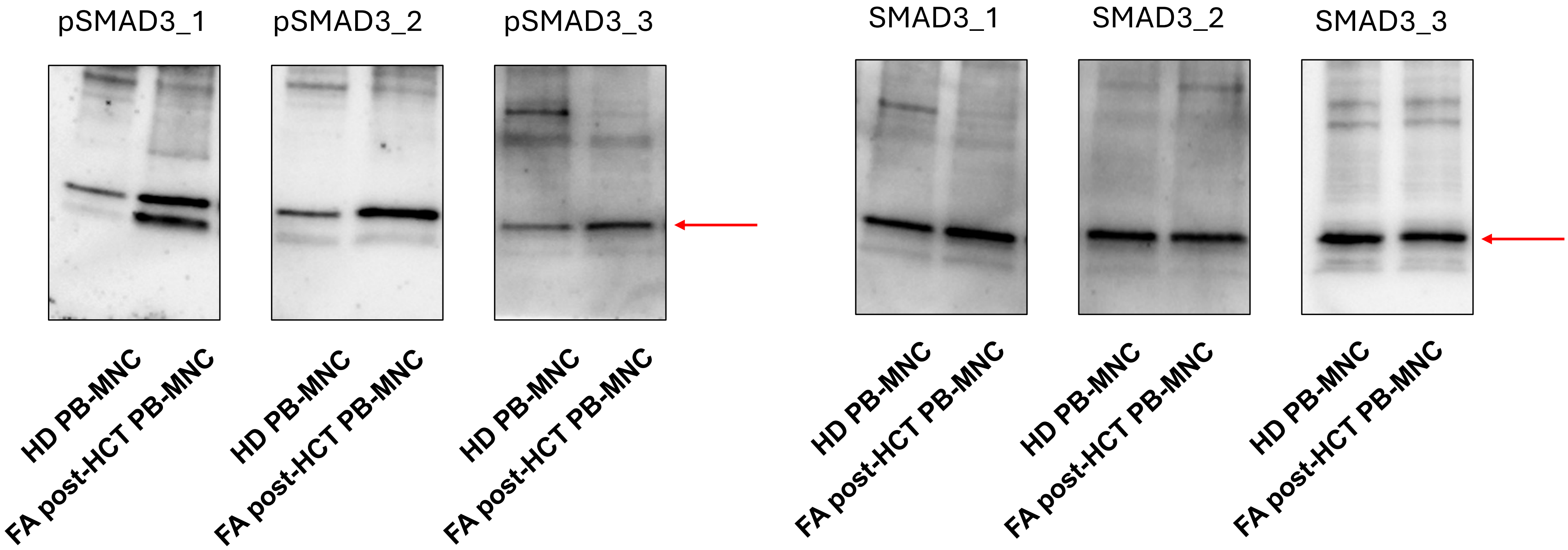

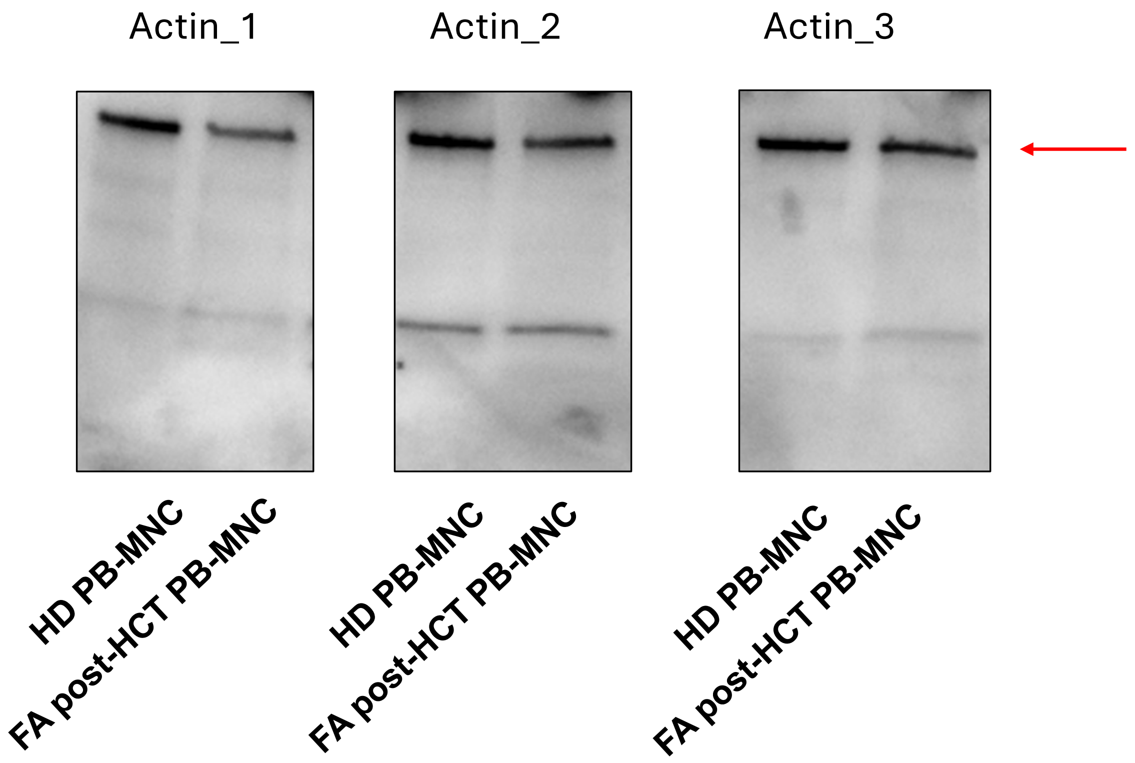
